# Dissecting epigenome dynamics in human immune cells upon viral and chemical exposure by multimodal single-cell profiling

**DOI:** 10.1101/2025.09.09.675101

**Authors:** Irem B. Gündüz, Bei Wei, Derek C. Chen, Wenliang Wang, Manoj Hariharan, Todd Norell, Timothy J Broderick, Micah T. McClain, Lisa L. Satterwhite, Thomas W. Burke, Elizabeth A. Petzold, Xiling Shen, Christopher W. Woods, Vance G. Fowler, Felicia Ruffin, Parinya Panuwet, Dana B. Barr, Aaron J Wilk, Madeline J Lee, Catherine Blish, Flora Castellino, Anna Maria Walley, Thomas Evans, Joseph R. Ecker, Fabian Müller, William J. Greenleaf

**Author notes:** Equal contribution (co-first) (Irem B. Gündüz, Bei Wei).

## Abstract

Environmental and pathogen exposure can lead to profound remodeling of the gene-regulatory landscape across human immune cell populations. Here, we present a single-nucleus chromatin accessibility atlas of human immune cells comprising 271,299 cells and 319,420 candidate regulatory elements from individuals exposed to HIV-1, COVID-19, Influenza virus, organophosphates, as well as healthy controls. Our longitudinal HIV cohort reveals epigenetic signatures of T cell exhaustion manifested in changes in the accessibility at binding sites for the FOXP family transcription factors. Upon severe exposure to SARS-Cov2 we identified changes in chromatin accessibility manifesting in a regulatory switch in cytokine networks characterized by the downregulation of the NF-κB motif family, alongside concordant changes in AP-1 and IRF factor networks in CD14+ classical monocytes. By integrating single-cell profiles of DNA methylation from matched samples, we created a multimodal epigenome atlas of human immune cell states using the accessibility-derived candidate regulatory elements. Both modalities exhibit complementary epigenetic signatures at transcription factor binding sites associated with cell state, as exemplified in the process of memory formation in T-cells, where BATF, AP-1, and ETS motifs exhibit significant epigenetic covariance across both epigenomic layers. Finally, by linking potentially regulatory DNA methylation signatures to changes in chromatin accessibility in monocytes, we observe that severe COVID-19 involves selective, multiomics remodeling of epigenetic profiles at TF binding sites manifested in concordant DNA methylation and accessibility dynamics at inflammation-associated regulatory TFs.

## Introduction

Recent advances in sequencing and single-cell technologies have allowed the detailed molecular characterization of exposure-associated changes in the gene expression of human immune cell populations, revealing potential markers for diagnosis and therapeutic interventions [1,2]. For example, single-cell RNA sequencing profiling of peripheral blood mononuclear cells (PBMCs) from patients exposed to Severe Acute Respiratory Syndrome Coronavirus 2 (SARS-CoV-2) highlighted downregulated expression of pro-inflammatory cytokines in peripheral monocytes associated with the severity phenotype [3]. In contrast, other viral exposures, such as the influenza virus, lead to upregulated cytokine expression in peripheral monocytes [2]. The expression patterns of cytokine receptor genes and transcription factors also differ in Influenza virus exposure from SARS-CoV-2 exposure [4]. Mild cases of COVID-19 exhibit a more limited activation of NK and monocyte cells compared to mild cases of influenza. [5]. Exposure to the Human Immunodeficiency Virus type 1 (HIV-1) induces changes in cytokine and chemokine expression, similar to SARS-CoV-2 exposure, leading to shifts in cell composition and increased signatures of inflammation [6]. However, while similarities and differences in molecular responses between COVID-19 and other viral exposures are well studied at the transcriptomic level, the underlying gene regulatory programs that contribute to these responses remain less well characterized. Exposure to chemicals, such as organophosphate pesticides (OP), can lead to toxic effects similar to those observed with exposure to infectious diseases [7]. For example, low-dose chronic OP exposure is associated with increased expression of pro-inflammatory cytokines, such as IL-1β [8]. Population-level reports link chronic OP exposure to differential methylation levels in cellular defense-associated genes, such as the HLA family [9]. However, the gene regulatory programs of human immune cells in response to OP exposure remain to be characterized at high resolution.

Single-nucleus epigenomic profiling-based analysis using the assay for transposase-accessible chromatin with sequencing (ATAC-Seq) [10] has been used to characterize gene regulatory programs of human peripheral blood cells in healthy individuals and across various disease states [5, 11, 12, 13]. A small number of studies with small cohort sizes have reported *in vivo* accessibility datasets of human immune cells upon exposure to infectious agents [14, 15]. However, a more comprehensive and integrative analysis of gene regulation across exposures to infectious agents as well as to chemicals could provide a more holistic view of common and distinct regulatory programs the immune system uses to respond to environmental stimuli.

To chart the regulatory changes conferred by dynamic chromatin accessibility, we performed snATAC-seq [10] on 92 samples collected from individuals exposed to SARS-CoV-2, HIV-1 virus, Influenza A virus (IAV), and organophosphate pesticide (OP) exposures, along with healthy donors, and we generated chromatin accessibility profiles of 271,299 peripheral blood cells. We analyzed how these exposure-specific alterations affect the activity of *cis-*regulatory elements (cCREs) in each exposure state and cell type. Individuals exposed to SARS-CoV-2 exhibited significantly altered chromatin accessibility profiles in CD14+ monocytes, linked to a potential shift in TF-based regulation of cytokine signaling networks, such as the downregulation of the NF-kB motif family. Furthermore, we identified chromatin signatures of T cell exhaustion in prolonged HIV-1 infection, characterized by changes in accessibility of regulatory networks involving FOXP transcription factors. By integrating single-cell methylation profiles from matching cohorts [16,17], we further characterized the interplay between DNA methylation and chromatin accessibility in potentially regulatory DNA regions and found anticorrelated patterns of chromatin accessibility and DNA methylation in these regions that indicate coordinated epigenetic changes across cell populations and epigenomic modalities. Our analysis revealed TF signatures such as BATF, AP-1, and ETS transcription factor (TF) motif families, exhibiting joined dynamics in accessibility and DNA methylation in memory T-cells compared to naive states, suggesting active regulatory roles in T-cell memory formation. Finally, we identified changes in DNA methylation of cCREs that are concordant with accessibility changes observed upon SARS-CoV-2 exposure in CD14+ monocytes, particularly in regulatory networks involving Interferon regulatory factors (IRFs) and AP-1 factors.

## Results

### A single-cell epigenome atlas of human immune cells across viral and chemical exposures

We assayed the single-cell chromatin accessibility landscape of human immune cells of healthy individuals and individuals exposed to viral and chemical agents. We profiled 92 human PBMC samples across 4 different cohorts and exposure groups (Figure 1A; Supplementary Table 1): (i) a SARS-Cov2 cohort consisting of healthy control samples (n=7) and individuals with mild (WHO score 1–3; n=8), moderate (WHO score 4-5; n=4) and severe (WHO score 6-8; n=6) COVID-19 indications; (ii) a longitudinal cohort of four HIV infected individuals in pre-infection (n=4), acute infection (n=4) and chronic infection (n=4), in which antiretroviral therapy (ART) was administered between the acute and chronic infection stages; (iii) an influenza cohort of 5 individuals sampled 2 days prior to live influenza virus challenge (control group) as well as 3 days, 6 days and 28 days after virus challenge; and (iv) a cohort of individuals with varying degrees of exposure to organophosphates (OP), grouped by urinary TCPY (3,5,6-trichloropyridinol) levels: low exposure (TCPY < level of detection; n=9), medium exposure (0 < TCPY <= 2.4 μg/L; n=6) and high exposure (TCPY > 2.4 μg/L; n=20).

**Figure 1:**
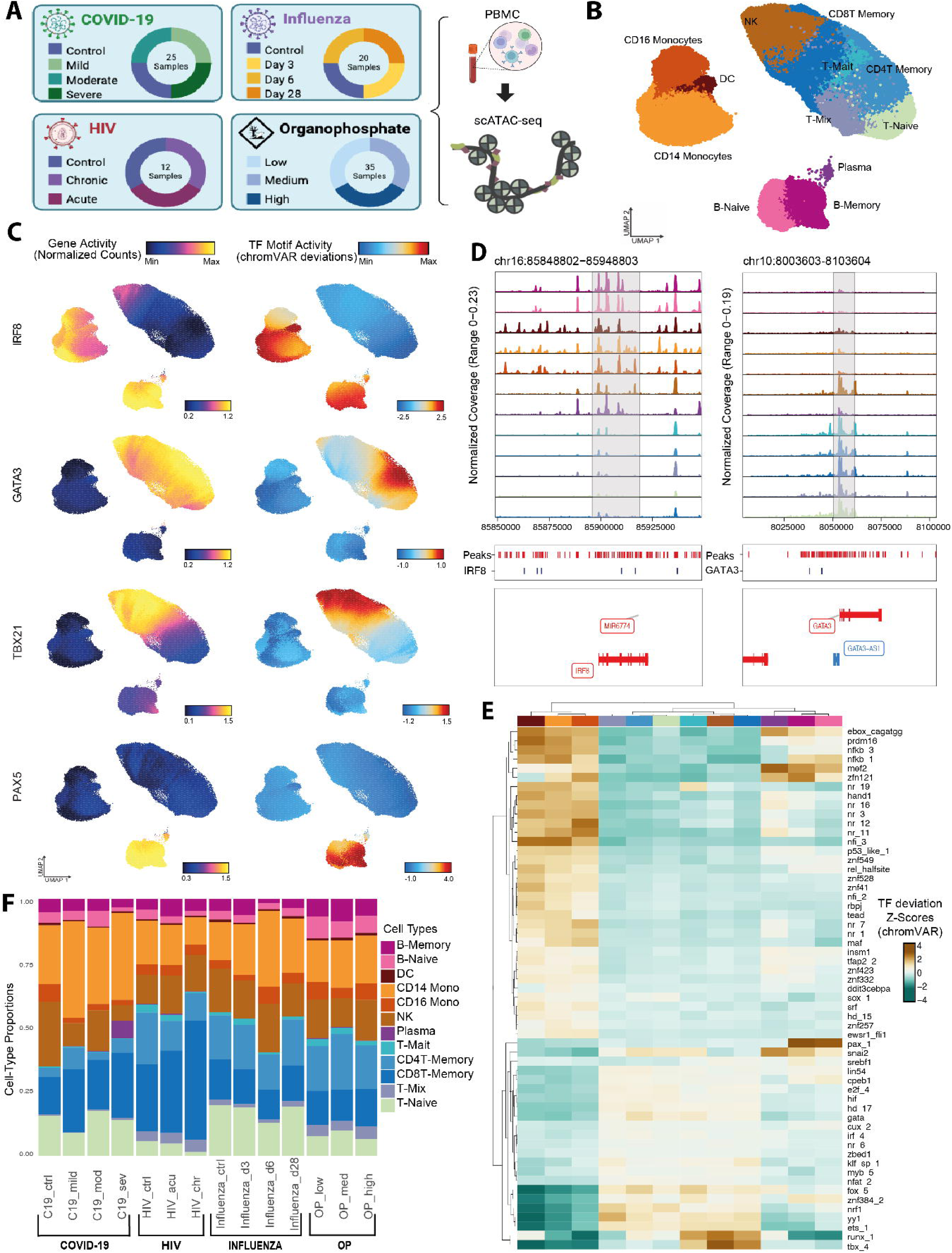
The chromatin accessibility landscape of human immune cells across viral and chemical exposures. A) Overview of sample cohorts and single-cell chromatin accessibility profiling B) UMAP dimensionality reduction of the 271,299 cells from the scATAC dataset colored by immune cell population C) UMAP dimensionality reduction colored by the relative gene activity scores (left) and transcription factor (TF) motif activity (ChromVAR, right) of IRF8, GATA3, TBX21, and PAX5 D) Genome accessibility tracks by cell type with a focus on genes (IRF8 and GATA3) and, their respective motif occurrences E) Heatmap of the 57 top variable TF motif cluster [45] deviation Z-scores (ChromVAR) across cell types F) Bar plot of cell type composition across cohorts and exposure groups

We performed a 10X-based single-nucleus assay of transposases accessible chromatin sequencing (snATAC-seq) and obtained chromatin accessibility profiles of 285,276 cells, of which 271,299 cells were retained after quality control and filtering (Supplementary Figure 1A; Methods). We integrated our four cohorts to characterize immune cell populations across pathogen exposures while correcting for batch effects using Harmony [18]. We identified 22 clusters using graph-based clustering (see Methods, Supplementary Figure 1B) and 319,420 reproducible peaks that represent candidate *cis*-regulatory elements (cCREs) across the identified clusters (Supplementary Figure 1C).

To annotate these clusters with corresponding cell types, we integrated and mapped cell type annotations from a publicly available single-cell transcriptome atlas of cross-tissue immune cells [19] (Supplementary Figure 1D). We grouped our 22 clusters into 12 broad immune cell populations, including NK cells, CD14^+^ classical and CD16^+^ non-classical monocytes, dendritic cells (DC), as well as naive and mature subpopulations of B cells, plasma cells, and T cells (Figure 1B). In the T-cell lineage, we combined several less-abundant or closely related subpopulations into a single group that we termed “mixed T-cells” (T_mix). This group includes CD4/CD8 T-memory cells, T follicular helper (Tfh), T gamma delta (Tgd) cells, regulatory T cells (Tregs), and specific subsets of resident memory T cells (Trm).

To validate these annotations, we examined the chromatin-derived gene activity and transcription factor (TF) activity, which serve as a proxy for gene expression and regulatory potential, of canonical cell type markers (Figure 1C, D, and Supplementary Figure 1E) [11]. Gene loci of corresponding marker genes exhibited characteristic chromatin accessibility profiles around promoter and cCRE regions in their respective cell types (Figure 1D and Supplementary Figure 1E, F), e.g., IRF8 exhibited increased activity in B cells and monocytes, while the GATA3 locus and its target binding sites were mostly accessible in T and NK cell populations.

To better understand the role of transcription factors in the regulation of cell state, we aggregated accessibility across the genomic occurrences of a consolidated set of clustered TF binding motifs (non-redundant archetype sequences created by grouping highly similar motifs, referred to as Altius motif set) (Figure 1E) [20,21]. Comparing the derived TF motif activities of cells with each other (Supplementary Table 2), we observed increased activity of NF-kB cluster (RELA, RELB, and NF-κB1/2) in CD14^+^ classical and CD16^+^ non-classical monocytes, dendritic cells (DC), B naive, B memory, and plasma cells compared to T and NK cell populations. We also observed increased chromatin activity at candidate binding sites of transcription factors known to regulate cellular differentiation and function in T cells, specificallyRUNX (RUNX1-3) and ETS (ETS1, ELF1) clusters, and B cells’ PAX cluster (representing PAX5), compared with other cell types.

Next, we assessed the changes in cell-type composition across exposures (Figure 1F). We focused on within-cohort comparisons of the cell-type composition in the exposed samples with the respective control groups (Supplementary Figure 1G). We observed a relative expansion of the CD8^+^ T cell population (padj=0.029) and a consistent but statistically insignificant depletion of CD4^+^ T cells in individuals chronically exposed to HIV (Supplementary Figure 1G). In individuals with severe COVID-19 conditions, we observe a proportionally larger population of plasma cells (padj=0.016) and depletion of NK cells (padj=0.035). Other cell types did not exhibit significant changes in proportions in severe cases of COVID-19, consistent with reports by Wilk 2021 [5]. Furthermore, we observed a depletion of naive T-cells (padj=0.04) and DC cells (padj=0.022) in mild cases of COVID-19 compared to the control group. Although the cell-type compositions do not change significantly during OP and influenza exposures, we observed limited composition changes. For example, on the 6th day after Influenza exposure, we observed a depletion in T-naive cell composition and an increase in classical CD14+ and non-classical CD16+ monocytes compared to controls (Figure 1F, Supplementary Figure 1G, padj = 0.056, 0.15, respectively).

To map the landscape of exposure-related changes across various cohorts and cell types, we also compared the TF activity signatures associated with each exposure group (Supplementary Figure 2A). Notably, comparing exposure groups to their respective control samples did not reveal strong shared exposure-induced changes in chromatin accessibility derived TF activities. Instead, we observe that TF motif activity was primarily defined by cell lineage identity, with cohort-specific signatures appearing as a secondary effect (Figure 1E, Supplementary Figure 2A). We identified a shared stress-response program driven by AP-1 factor motifs (ap_1; FOS, JUN, FOSL1) that was selectively activated in myeloid subsets (CD14+ monocytes and dendritic cells (DC)) across all exposure cohorts. In contrast, non-classical monocytes (CD16+) and lymphoid cells maintained lower accessibility across those TFs. Furthermore, B-cell populations (naive and memory) were characterized by high NF-kB activity (nfkb_1/REL, NF-κB1) rather than AP-1, while T and NK cells showed little to no activation of these factors. Beyond these shared responses, we observe TF activity signatures specific to exposure cohorts. For instance, the higher interferon response (irf_1/IRF1) accessibility was restricted to OP and HIV exposures within NK and T-cell populations compared to other exposures (COVID-19, Influenza), and HIV and OP exposure cohorts showed similarity by exhibiting lower AP-1 motif (ap_1; FOS, JUN, FOSL1) activity in memory and naive T-cells.

### HIV-induced T-cell exhaustion dynamics in different stages of the infection

To further characterize the expansion of the CD8^+^ T cell population during HIV infection, we sub-clustered CD8^+^ T cells into different subtypes. Of note, we identified a population of exhausted T cells (Tex), marked by higher gene activity scores (defined as aggregated locus accessibility, see Methods) of exhaustion markers such as CTLA4, NCAM1, and HAVCR2 (Figure 2A-C, Supplementary Table 3) [22]. We also found other gene activity scores elevated in this subset, such as ROBO1/2, which are not traditional exhaustion markers, suggesting specific roles of these genes in HIV-infection-related exhausted T cells [23,24].

**Figure 2.**
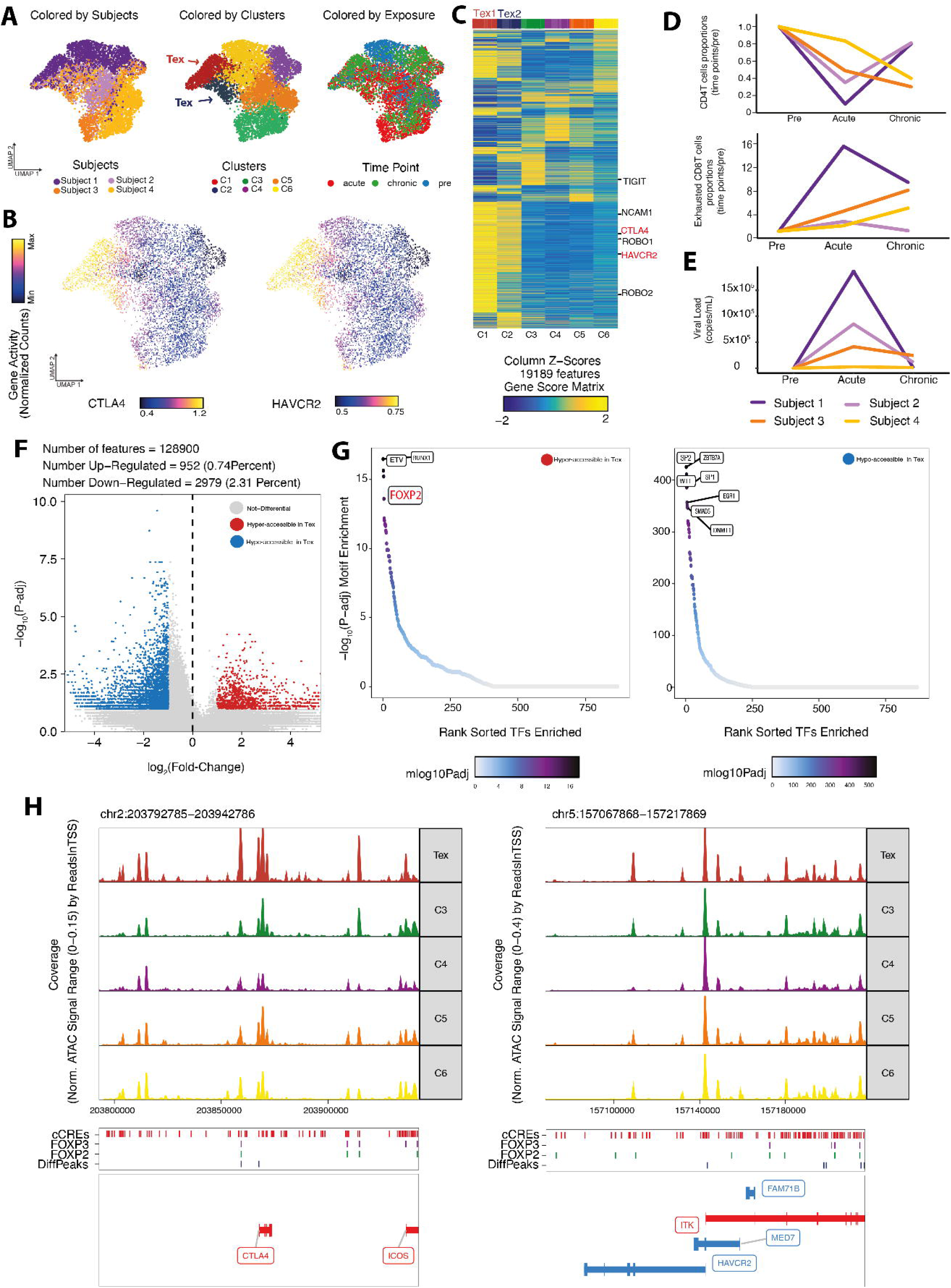
Chromatin accessibility dynamics in T cell populations upon HIV exposure. A) UMAP projection of the T-cell subpopulation from the HIV-subsetted scATAC dataset colored by subjects (first), colored by Louvain clusters (middle) and, colored by time point (last) B) Side-by-side UMAP projections of the HIV-subsetted scATAC dataset colored by gene activity scores of T-cell exhaustion markers CTLA4 and HAVCR2 C) Heatmap representation of differentially accessible marker genes in different clusters in CD8+ T cell population based on gene activity scores D) Cell type proportion relative to the pre-stage over time for exhausted CD4+T cells (upper) and exhausted CD8+T cells (lower) per subject E) Viral load per subject over the disease progress of HIV F) Fold-change versus p-adjusted (Volcano) plot of exhausted (cluster 1 & 2) and non-exhausted T-cells in the HIV cohort, hyper-accessible differential peaks highlighted in red, hypo-accessible peaks highlighted in blue. Differential accessibility was defined using an FDR ≤ 0.01 and |log_₂_fold change| ≥ 1.25. G) TF motif enrichment plots in hyper-accessible (left) and hypo-accessible (right) H) Genome accessibility tracks per CD8 T subpopulation with a focus on T-cell exhaustion markers CTLA4 and HAVCR2, along with occurrences of the FOXP2 and FOXP3 motifs

When we assessed the proportional changes in T-cell populations within the HIV cohort, we observed a depletion of CD4^+^ T cells. In contrast, the proportion of exhausted CD8^+^ T cells increased after the HIV exposure (“acute” stage, Figure 2D). We confirmed a stepwise progression of exhaustion by comparing the proportion of Tex cells across infection stages. A significant accumulation of Tex cells began during the acute phase relative to the baseline (Fisher’s exact test, acute vs pre, p-value = 2.57e-28) and further increased during the chronic phase (Fisher’s exact test, acute vs chronic, p-value = 1.09e-10) (Figure 2A, UMAP colored by time point). Changes in cell abundance for CD4^+^ T cells and exhausted CD8+ T cells are anti-correlated throughout the progression of infection. For example, subject 1 exhibited the most severe depletion of CD4^+^ T cells along with an increase of exhausted CD8^+^ T cells during the infection trajectory; for subjects 1 and 2, the proportions of CD4^+^ T cells decreased in the “acute” stage and recovered in “chronic” stage after antiretroviral therapy (ART), consistent with a response in viral load to ART and regeneration of the CD4^+^ compartment (Figure 2E). Conversely, the proportions of exhausted CD8^+^ T cells decreased in the “chronic” stage compared to the “acute” stage; For subjects 3 and 4, the proportions of CD4^+^ T cells continued to decrease in the “chronic” stage after ART, while the proportions of exhausted CD8^+^ T cells also increased, highlighting that cell type compositions relevant to disease progression and therapeutic response can be inferred from epigenetic signatures.

Next, we identified differentially accessible regions in exhausted T-cells (Tex) compared to other CD8^+^ T-cell populations in the HIV cohort (Figure 2F, Supplementary Table 4). We then determined the activity changes of TF motifs based on the enrichment of corresponding motifs in hypo- and hyper-accessible cCREs. In regions with increased accessibility in exhausted cells, we observed the enrichment of FOXP2 and other FOX family motifs, while elements with decreased accessibility were enriched in SP and SMAD motifs (Figure 2G). We then examined the TF motifs present around accessible elements near marker genes for T cell exhaustion; for example, for gene CTLA4, we found that FOXP motifs are located mostly within cCREs with increased accessibility in Tex compared with other subtypes/subclusters of CD8^+^ T cells (Figure 2H), consistent with the hypothesis that CTLA4 was activated by FOXP family binding in Tex.

### SARS-CoV-2 Reshapes Chromatin and Cytokine Regulation in CD14^⁺^ Monocytes

To assess the impact of each exposure on cell-type-specific accessibility profiles, we aggregated the accessibility of each cell type for donors and exposure conditions into pseudo-bulk samples. We performed pairwise differential accessibility analysis between control and disease groups for each cohort and cell type across cCREs. Our analysis revealed limited differentially accessible regions (DARs) upon influenza and organophosphate exposures (Supplementary Figure 3A). In contrast, we observe substantial differences in accessibility associated with COVID-19, particularly in severe cases within the population of CD14^+^ classical monocytes (Figure 3A, Supplementary Figure 5A).

**Figure 3.**
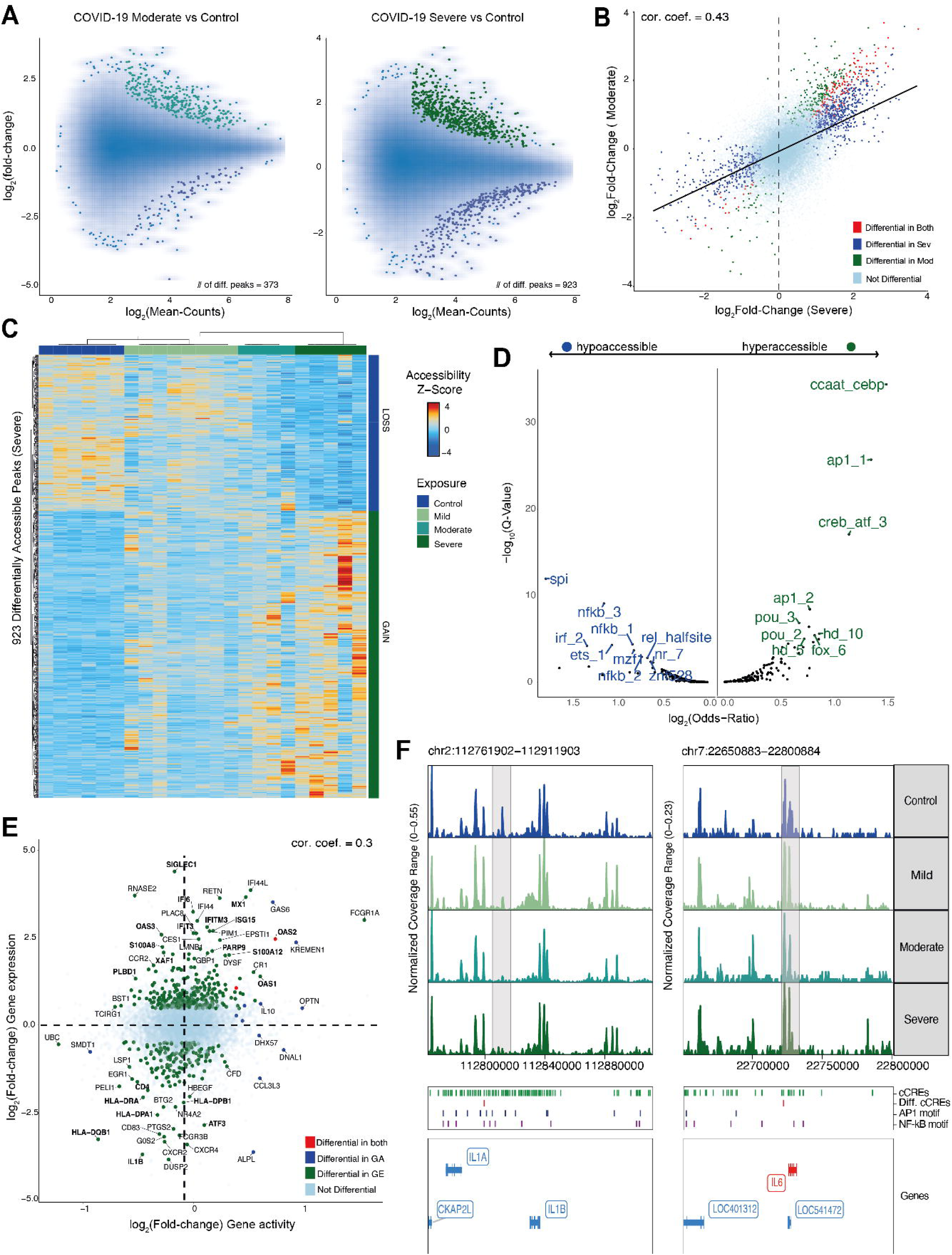
Impact of COVID-19 on chromatin accessibility profiles of CD14+ classical monocytes. A) Fold-change versus mean count (MA) plots comparing moderate (left) and severe (right) cases in the COVID-19 cohort to healthy controls, hyper- and hypo-accessible DARs (p.adj < 0.05, log2fold-change > 0.5) are highlighted in green and blue, respectively. Point density is visualized as a 2D gradient; low-density outliers are shown in addition (< 0.05, | log_₂_(fold-change) >= 0.5|). B) Fold-change in severe COVID-19 cases vs healthy control compared to fold-change in moderate cases in CD14^+^ classical monocytes with fitted regression line. All peaks are shown; DARs are highlighted as in A). C) Heatmap of Z-score normalized accessibility profiles of 923 DARs (p.adj < 0.05, |log=(fold-change) >= 0.5|) in CD14^+^ classical monocytes in mild, moderate, and severe cases of COVID-19, along with control samples. Hierarchical clustering with Euclidean distance (complete linkage) was applied to both rows and columns of the heatmap. D) Enrichment analysis of TF motif clusters in DARs (Fisher’s Exact Test using LOLA) upon severe COVID-19 exposure compared to control [45]. Enrichment in hyperaccessible cCREs is highlighted in red, while enrichment in hypoaccessible cCREs is highlighted in blue. E) Scatterplot comparing the fold-change in differential gene activity (derived from chromatin accessibility) versus the fold-change obtained from differential gene expression (DEG) analysis, with genes determined as DEGs from Wimmers 2023 and Wilk 2021 highlighted [26,5]. F) Genome accessibility track visualization of the differentially accessible cCREs at the IL1B and IL6 gene loci. cCREs and TF motif occurrences are annotated as separate tracks.

To systematically assess the coherence of differences in chromatin accessibility across exposures, we computed pairwise correlation coefficients of fold-changes of exposure groups to corresponding control groups of cells (Supplementary Figure 4). This analysis revealed the differences in accessibility dynamics in viral and chemical exposures, manifested in distinct correlation blocks for each exposure cohort. While COVID-19 induced significant changes distinct from other viral exposures, we also observed a weak but notable positive correlation between COVID-19 and Influenza induced changes in chromatin accessibility, specifically within the CD14 monocyte population. This suggests a potential shared chromatin signature between these respiratory viruses that is restricted to innate immune response rather than being a global feature of the antiviral response.

We identified 923 differentially accessible regions (DARs) in CD14+ classical monocyte profiles in severe COVID-19 and 373 DARs in moderate conditions compared to control conditions (Figure 3A). We found that in the union of these DARS, accessibility fold-changes positively correlate between moderate and severe conditions, indicating that these underlying differential regions are consistent across these levels of disease severity, differing primarily in the magnitude of the observed accessibility changes (Figure 3B, C). Next, we performed TF motif enrichment analysis in DARs, revealing an enrichment of hypoaccessible regions in severe cases of COVID-19 for NF-kB and SPI motifs. In contrast, hyperaccessible regions were enriched for AP-1 and CREB motifs (Figure 3D). Both NF-kB and AP-1 are key regulators of cytokine signaling; however, while NF-kB is predominantly associated with acute inflammatory responses, AP-1 factors are linked to late-stage inflammation and stress responses. A shift from NF-kB to AP-1-dominated regulation may reflect an underlying immune paralysis, consistent with previous reports (Wilk 2021, Wimmers 2023, and Lam 2023 [25,26].

To evaluate how chromatin accessibility changes relate to gene expression changes, we re-analyzed publicly available, matched single-cell transcriptome data from 34,548 cells obtained from the same COVID-19 cohort (see Methods) [3]. We compared differential gene expression in CD14^+^ classical monocytes between severe COVID-19 and control groups from scRNA-seq and gene activity derived from chromatin accessibility (Figure 3E, Supplementary Table 5, see Methods). Consistent with the findings of Wimmers 2023 [26], we observed an increased expression and gene activity in interferon-stimulated genes (ISGs), including the IFIT family (Figure 3E, Supplementary Figure 5B). In contrast, concordant decreased expression and gene activity were observed in primarily inflammatory genes, including IL-1B. We also identified genes proximal to cCREs with decreased accessibility that align with the findings of Wilk 2021, such as IL6 and CD4 (Figure 3F, Supplementary Figure 5B). We observed decreased accessibility in CD4, a key regulator of macrophage differentiation and pro-inflammatory cytokine induction, under severe conditions (Supplementary Figure 4B) [5,27].

An enrichment analysis of gene function linked to cCREs gaining accessibility revealed an enrichment in pathways governing apoptosis and the negative regulation of type I interferon-mediated signaling, suggesting a potential shift toward immune suppressive and pro-apoptotic programs (Supplementary Figure 5C). This global chromatin reprogramming included the targeted silencing of genes that define classical monocyte identity. Specifically, we observed decreased accessibility at the CD4 locus, which contains RELA binding motifs (Supplementary Figure 5B). Consistent with global hypoaccessibility of NF-kB motifs in severe cases (Figure 3D), this suggests that the implied decrease in RELA binding contributes to the silencing of these genes. Consequently, we observed a significant and coordinated decrease in accessibility and expression at several HLA Class II genes, specifically HLA-DRA, HLA-DPA1, and HLA-DQB1 (Figure 3E, Supplementary Figure 5B). The silencing of these loci, which are key components of “MHC class II protein assembly” and “antigen presentation” pathways, suggests that CD14+ monocytes in severe COVID-19 adopted MDSC-like molecular features [5,28]. Together, these findings reveal a chromatin reprogramming pattern in severe disease, where the opening of motifs driving immune suppression and the closing of antigen presentation genes act as primary drivers of altered monocyte function.

### Complementary Chromatin and DNA Methylation Signatures of Immune Cell States

We integrated our single-cell chromatin accessibility atlas with available single-cell DNA methylation data, which were separately generated using snmC-seq2 for a subset of 39 samples from our organophosphate and HIV cohorts using snmC-seq2 [16]. This methylome atlas comprises single-cell profiles for 7 major immune cell types obtained by fluorescence-activated cell sorting (FACS) (see Methods). We aggregated DNA methylation signals across the cCREs identified by scATAC-seq. After preprocessing and quality control (Methods), we retained 22,488 cells with DNA methylation profiles, which we integrated with corresponding chromatin accessibility profiles of 110,376 cells from matching samples using canonical correlation analysis (Methods) (Figure 4A). This integration on the level of cCREs allowed the cross-projection between low-dimensional representations of chromatin accessibility and DNA methylation and the transfer of annotated cell types between modalities. We observe high cell-type correspondence between the annotated chromatin accessibility and DNA methylation profiles on the level of broadly defined cell types (B-cells, T cells, monocytes, NK cells) while noting weaker correspondence on the level of T-cell subpopulations (Figure 4A, B) and the individual cell level.

**Figure 4.**
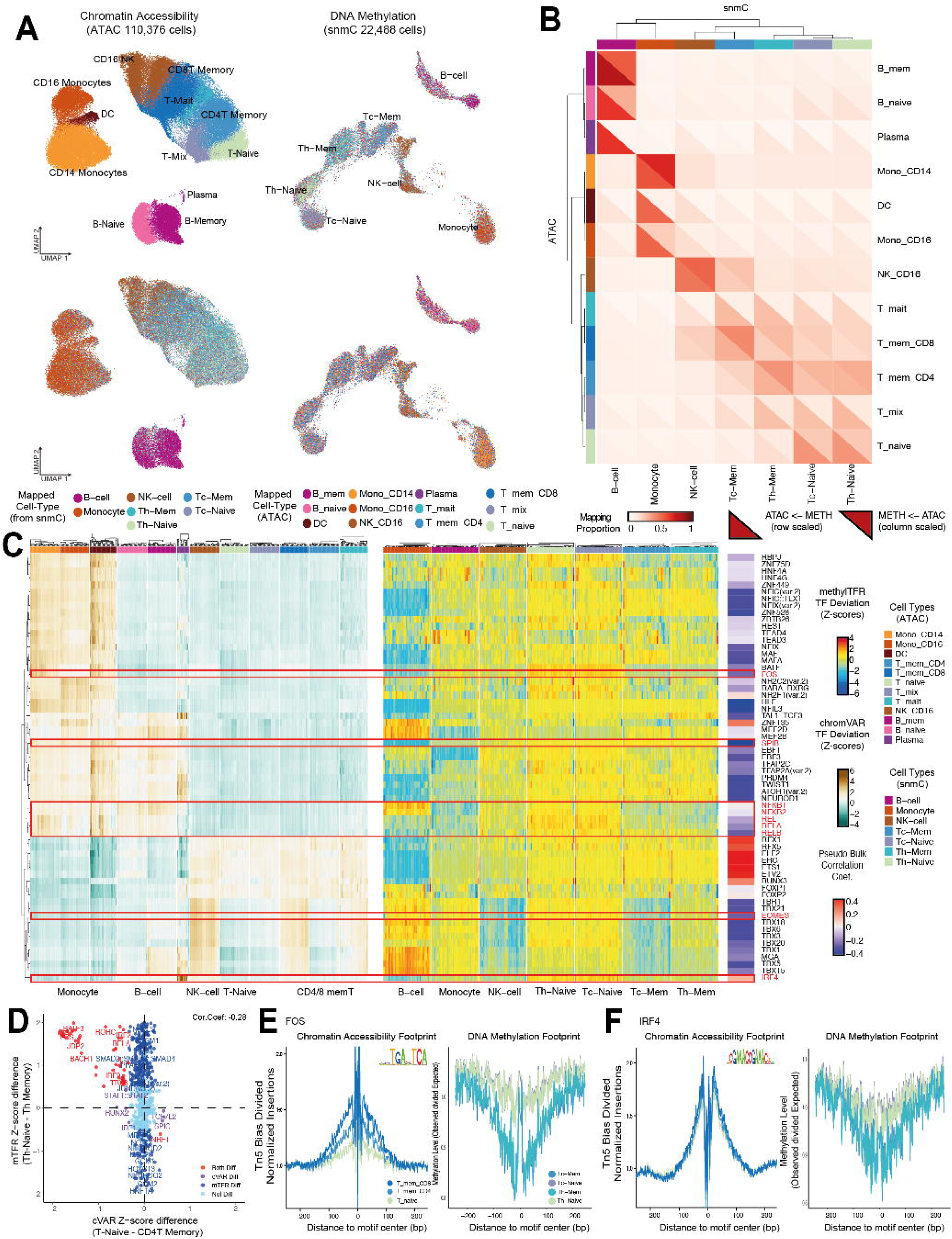
Integration of chromatin accessibility and DNA methylation, epigenetic TF activity across cell states. A) UMAP projections of 110,376 cells from scATAC with cluster cell labels (top-left) and mapped snmC labels (bottom-left), UMAP projection of 22,488 cells based on DNA methylation im cCREs with FACS sorted cell labels (top-right) and mapped ATAC labels (bottom-right). B) Heatmap of pairwise cell-type correspondence between scATAC-seq and snmC-seq. Upper triangles: projection of scATAC labeling onto the snmC manifold; lower triangles: snmC labeling onto the scATAC manifold. Correspondence is denoted by the proportion of mapped cells per input cell type. C) Heatmap of 61 most variable motifs with their chromVAR deviation Z-scores (left) and their corresponding methylTFR deviation Z-scores (right), along with row-wise Pearson correlation coefficients calculated between the chromVAR and methylTFR deviation scores across all pseudobulk populations and methyl-SELEX [29] TF annotation. D) TF motif activity Z-score difference (CD4T-Memory Z-score subtracted from T-naive Z-score) between chromatin accessibility and DNA methylation-based TF activities. The overall Pearson correlation is -0.28, and differentially active motifs identified from a two-sided Wilcoxon test (padj < 0.05) are highlighted. E) Side-by-side accessibility and methylation footprint plots of FOS and F) IRF4 motifs in naïve and memory T cell populations.

Global transcription factor (TF) activity, when inferred from chromatin accessibility variability at targeted motifs, is often strongly correlated with epigenetic changes of those sites [21]. The role of DNA methylation in TF binding sites is complex: although methylation can promote TF binding in some motifs, it can prevent binding in others [29]. The mechanism underlying the interaction between DNA methylation and chromatin accessibility in regulating TF activity is not yet fully understood. While our integration demonstrated global correspondence between accessibility and methylation profiles at the cCRE level, we also examined the combined effects of the two modalities at the TF binding site (TFBS) level.

To determine the relationship between chromatin accessibility and DNA methylation at regulatory DNA elements, we derived TF regulator activity by aggregating chromatin accessibility [21] and DNA methylation profiles across putative binding sites of TFs in pseudobulk samples for each individual, disease condition, and cell type (see Methods and Supplementary Figure 6A). We observed that TF activities derived from both epigenomic modalities can discriminate broad cell type clusters, with methylation-based TF activities additionally providing fine-grained differentiation between naive and memory T-cell clusters (Supplementary Figure 6B).

We identified TF motifs exhibiting the highest variability in chromatin accessibility-based TF activities across cell types. We observed a moderate inverse relationship between accessibility and methylation-based TF activity scores across cell types, where TFs exhibiting higher accessibility tended to show lower methylation and vice versa (Figure 4C). Analyzing this relationship across cell types, we observe that this inverse correlation was most pronounced in B-cells (cor.=-0.48), followed by Th-Naive (cor.=-0.23), Monocyte (cor.=-0.18), Tc-Naive (cor.=-0.17), NK-cell (cor.=-0.17), Th-Mem (cor.=-0.01), and Tc-Mem (cor.=0.01), respectively (Supplementary Figure 6D, Supplementary Table 6). For instance, the SPIB motif displays high accessibility with low methylation in monocytes, whereas in T-cells, it exhibits lower accessibility accompanied by higher methylation. A similar inverse relationship between accessibility and methylation was observed for NF-κB family members, including REL, RELA, and RELB (Figure 4C, Supplementary Figure 6C).

To further assess the methylation sensitivity of these motifs, we examined their annotated methylation sensitivity from published methyl-SELEX data [30,31]; 397 JASPAR motifs correspond to TFs annotated with methylation sensitivity annotation comprising evidence of negative impact of methylation on binding (methylMinus), and positive effects (methylPlus). We observed a broad range of correlation scores between accessibility and methylation-associated TF motif activity for methylMinus and methylPlus motifs, with methylMinus motifs exhibiting a skew towards negative correlation coefficients between methylation and accessibility (Supplementary Figure 6D).

Consistent with reports of global loss of DNA methylation in T cell memory formation [33], we observe lower methylation levels at TF motifs in memory T cells compared to naive cell states (Figure 4C, D). We observed that naive and memory T-cells exhibit variable methylation activity patterns across TF motifs, accompanied by relatively stable accessibility patterns. To characterize the interplay between methylation and accessibility in TF-driven regulation of T-cell memory formation, we compared motif activities between naive and memory T-cells across both modalities (see Methods). We observed an anti-correlation between the differentially active motifs identified from the two epigenomic modalities in CD4^+^T and CD8^+^T cell subsets, suggesting that these epigenetic layers coordinate TF activity to drive memory formation in T-cells (Figure 4D and Supplementary Figure 7A).

We assessed aggregate chromatin footprints – characteristic high accessibility at TFBS compared to the genomic background and protective patterns likely due to TF binding – of TF motifs implicated with roles in T-cell memory formation. We compared these aggregate chromatin accessibility footprints to DNA methylation footprints (Figure 4E, F, Supplementary Figure 7B). Our findings indicate that BATF, FOSL2::JUND, and FOS motifs exhibit increased accessibility and decreased methylation levels in memory T-cells compared with naive cells, consistent with the accessibility reports of Calderon 2019 and methylation reports of Durek 2016 [32,33]. Furthermore, RELA, IRF4, and RUNX3 exhibit a limited increase in accessibility while exhibiting markedly decreased methylation in T-memory cells compared to naive T-cells. Consistent with observations from Abdelsamed 2020, who showed decreased methylation levels in the EOMES binding sites in CD8 memory T-cells [34], we also observe a decrease in methylation in T-memory cells compared to T-naive cells across EOMES binding sites, coupled with increasing accessibility to CD8^+^ T-memory cells compared to naive T-cells (Supplementary Figure 7B). MAF::NFE2 motifs show increased accessibility in memory T-cells compared to naive T-cells, with decreased memory-specific methylation observed (Supplementary Figure 6B). FOXP1 motifs, a known marker of memory formation in T-cells, exhibit slightly decreased methylation in memory T-cells, with limited decreased accessibility patterns in quiescent memory T-cells compared to naive T-cells [33]. The binding motifs of ETS1, a member of the ETS factor family that regulates T-cell differentiation and function, exhibit increased accessibility and decreased methylation in memory T-cells, compared to naive T-cells [35]. In contrast, NRF1 motifs displayed an inverse pattern, characterized by decreased chromatin accessibility and increased methylation levels in memory T-cells compared to naive states (Supplementary Figure 7A, B). Our findings thus suggest that memory-associated TFs are regulated by coordinated epigenetic changes at their binding sites, highlighting a layered regulatory program underlying memory formation in T-cells.

### Exposure to SARS-CoV-2 results in remodeling of NF-κB and AP-1 mediated regulation in CD14^⁺^ Monocytes

We next aimed to identify DNA methylation signatures that are associated with exposure-induced chromatin accessibility changes. We aggregated DNA methylation signals in cCREs defined by scATAC peak regions into pseudo-bulk samples per donor and exposure condition for each cell type and identified differential DNA methylation in cCREs by comparing healthy control and COVID-19 pseudo-bulk samples. Our results reveal that CD14+ classical monocytes exhibit the most pronounced differentially methylated regions (DMRs, i.e., differentially methylated cCREs) in severe cases of COVID-19 compared to other cell types (Figure 5A, Supplementary Figure 8A). In contrast, mild cases exhibit fewer DMRs, and other cell types show limited changes upon SARS-CoV-2 exposure. We observed 2073 DMRs in monocyte profiles in severe COVID-19 cases compared to healthy controls. We found a weak negative correlation (correlation coefficient = -0.07) between mean methylation difference and accessibility fold-change, indicating concordant epigenetic alterations across these modalities (Figure 5B).

**Figure 5.**
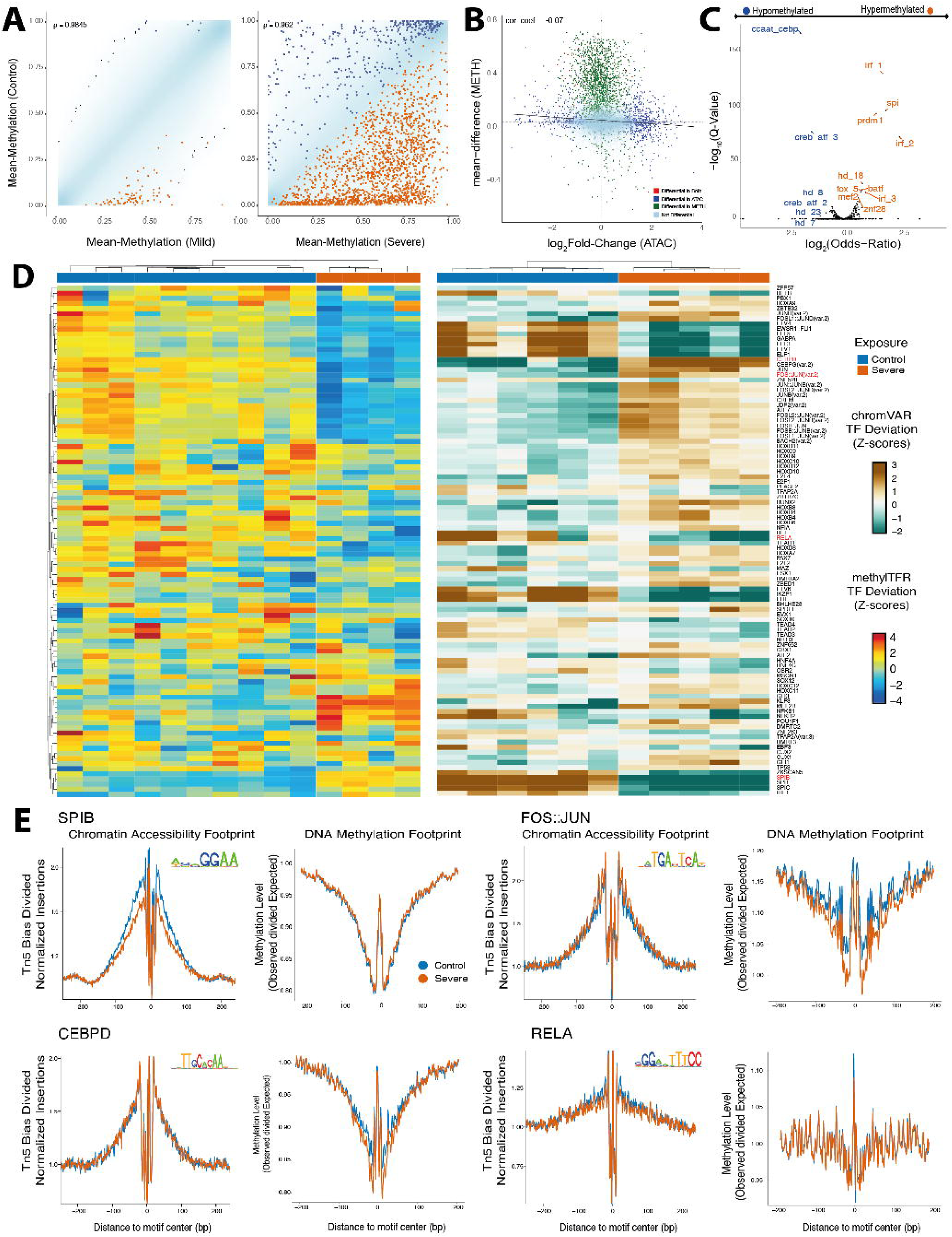
Impact of COVID-19 on DNA methylation profiles of cCREs in CD14+ classical Monocytes. A) Density scatter plots comparing aggregated DNA methylation levels in cCREs between commercial controls and mild (left, number of DMRs = 87) or severe (right, number of DMRs = 2073) COVID-19 samples. cCREs that are hypermethylated in the control group (adj. p < 0.05) are highlighted in blue, and cCREs hypermethylated in the severe group are highlighted in orange. Point density is visualized using 2D kernel density estimation; outliers in low-density regions (<0.1% of all points) are shown individually. B) Scatter plot of fold-change obtained from differential accessibility analysis between COVID-19 Severe and control samples versus the mean DNA methylation level difference in cCREs between COVID-19 Severe and control samples, including a regression line based on all cCREs. C) Enrichment analysis of TF motif clusters (LOLA) in hypermethylated and hypomethylated DMRs upon severe COVID-19 compared to control [45]. The enrichment in hypermethylated DMRs is highlighted in orange, while enrichment in hypomethylated DMRs is highlighted in blue. D) Heatmap showing row-wise Z-scores of differential transcription factor (TF) activity from control versus severe comparison: DNA methylation-based TF activity Z-scores (left) and chromatin accessibility-based differential TF activity Z-scores (right). Rows are clustered based on methylation profiles. E) Side by-side accessibility and methylation footprint plots of motifs with differential TF activity (SPIB, FOSL2::JUNB(var.2), CEBPD, RELA, respectively) (SPIB, FOSL2::JUNB(var.2), CEBPD, RELA, respectively)

Next, we performed TF enrichment analysis in the hypomethylated and hypermethylated cCREs of CD14+ classical monocytes in severe cases compared to controls, revealing enrichment of IRF1 motifs in hypermethylated cCREs consistent with reported expression changes from Wilk 2021 (Figure 5C) [5,36] and chromatin accessibility changes in these regions (Figure 3D). We observed the enrichment of IRF factors in hypermethylated DMRs consistent with methylation-based TF enrichment findings of Godoy=Tena 2022 [36]. We also found enrichment of CREB/ATF motifs in hypomethylated DMRs, which have been shown to inhibit the NF-κB-driven pro-inflammatory cytokine expression [37,38]. Consistent with the findings of Godoy=Tena 2022, we found enrichment of CEBP motifs in hypomethylated DMRs [36]. Moreover, consistent with the findings of Wimmers 2023 showing altered chromatin accessibility, we found enrichment of SPI motifs in hypermethylated DMRs [26].

To further characterize how the binding of these TFs could be altered through the epigenome upon SARS-CoV-2 exposure, we compared the epigenetic TF activity (pad.j < 0.05) based on chromatin accessibility and DNA methylation profiles at target binding sites (Figure 5D, Supplementary Figure 8B). We found that SPI factors lose accessibility and gain methylation in patients with severe COVID-19 compared to the control group. Moreover, AP-1 factors, such as FOSL2::JUN and JUN, gain accessibility accompanied by decreased methylation in severe conditions. In contrast, while NF-κB factors lose accessibility, we found that RELA and REL also lose methylation, whereas NF-κB1 and NF-κB2 gain methylation under severe conditions (Figure 5D).

Finally, we analyzed the motif footprints of differentially active TF motifs (Figure 5E, Supplementary Figure 8C). SPI motifs (SPIB and SPIC) exhibited reduced accessibility, while methylation levels for SPIB and SPIC showed marginal methylation differences from those of the control. This concordance was more pronounced when considering quantifications of TF activity based on aggregate signal in candidate binding sites (Figure 5D). Inflammatory AP-1 heterodimers (FOS::JUN) displayed a selective hypermethylation despite maintaining open chromatin. We observed slightly decreased accessibility in RELA and REL, NF-kB factors in the severe group, accompanied by unchanged methylation levels. For CREB and CEBP factors, such as CREB1, CEBPD, and CEBPA, we observed slightly increased accessibility and a limited decrease in methylation compared to the control group (Figure 5E, Supplementary Figure 8C). This effect was more apparent when considering quantitative scores of methylation and accessibility-derived TF activity (Figure 5D, Supplementary Figure 8C). Collectively, these patterns suggest a dysregulated state where monocytes repress their standard lineage identity (SPI) and specific inflammatory pathways (NF-kB), while simultaneously activating stress-response pathways (CEBP) and epigenetically constraining other inflammatory drivers (AP-1).

## Discussion

We profiled the single-cell chromatin accessibility of human immune cells across viral and chemical exposures. By high-resolution mapping of chromatin accessibility of candidate *cis-*regulatory elements (cCREs), we identified shared and stimulus-specific activity patterns that shape immune cell function and population differences in response to external stimuli. Furthermore, by integrating exposure-associated changes in chromatin accessibility with matched single-cell DNA methylation profiles, we characterized the epigenetic patterns associated with transcription factors (TFs) involved in defining cell type identity and during immune memory formation. In the epigenetic signatures that exhibited dynamics upon pathogen exposure, we observed the most striking differences upon SARS-CoV-2 exposure in CD14+ classical monocytes, and in this context, we further identified altered signatures of multimodal epigenetic activity of TFBS, including NF-kB and AP-1 factors. These findings highlight the utility of our single immune cell epigenome atlas for identifying regulatory mechanisms modulated by TF activity across cell states and in response to pathogens.

We identified an association between FOXP gene activity derived from accessibility and an associated enrichment of its potential binding motifs and the upregulation of exhaustion markers, particularly CTLA4, within a subset of CD8+ T cells from HIV-infected individuals. This parallels the well-characterized role of FOXP3 in CD4+ regulatory T cells (Tregs), where FOXP3 binding directly activates CTLA4 transcription to enforce immunosuppressive functions [54,55]. In the context of CD8+ T cell exhaustion, our data suggest a conserved regulatory mechanism wherein FOXP family members may drive CTLA4 expression to reinforce an exhausted phenotype. Although FOXP3 is not typically expressed at high levels in CD8+ T cells, the observed accessibility at FOXP motifs suggests that other FOXP family members may be active in this context. However, our current dataset lacks the protein or gene expression profiles to pinpoint the specific FOXP member(s) involved. Future studies employing single-cell transcriptome and TF binding profiling in HIV-specific CD8+ T cells could clarify this mechanism.

We found that classical CD14+ monocytes exhibit markedly altered chromatin accessibility at cCREs in COVID-19 and that the magnitude of these changes was correlated with disease severity. Target binding motifs of NF-kB factors exhibited decreased chromatin accessibility in severe COVID-19 cases, and we observed an enrichment of CREB/ATF factors among hyperaccessible cCREs, potentially contributing to the inhibition of NF-kB activity, consistent with previous reports [5,26]. We also found an enrichment of AP-1 factors in cCREs gaining accessibility in severe COVID-19, suggesting a shift in NF-kB-mediated regulation of the pro-inflammatory cytokine network to AP-1-mediated regulation. Previous studies indicate that silencing NF-kB factors turns CD14+ monocytes into a cell state resembling myeloid-derived suppressor cells (MDSCs) during severe COVID-19, resulting in decreased expression of HLA class II genes [5]. In line with this observation, we found decreased accessibility in the HLA class II gene vicinity.

Our integration of aggregated DNA methylation levels in cCREs with scATAC-seq data in paired cohorts demonstrated an anticorrespondence between these modalities at the broad cell-type level. This anticorrespondence was less pronounced when only considering T cell subpopulations of cells, indicating complementary information content of both epigenetic modalities. This discrepancy suggests the transition from naive to memory state in T-cells may involve regulatory mechanisms captured indirectly in chromatin accessibility, with DNA methylation providing an additional layer of information defining these cell states. DNA methylation and chromatin accessibility may be dynamic at different rates during the process of memory formation. Specifically, methylation-sensitive transcription factors (TFs), as revealed by methyl-SELEX [30], can exhibit distinct binding behaviors; DNA methylation suppresses TF binding in methylMinus motifs, whereas DNA methylation promotes TF binding in methylPlus motifs. Moreover, we developed a method to infer transcription factor (TF) activity from the DNA methylation footprints in TFBS, providing a complementary perspective on accessibility-based measurements of TF binding dynamics. We then systematically compared TF activities inferred from DNA methylation with those derived from accessibility. Our evaluation revealed that while accessibility-based TF activity provides a robust method to distinguish T-cell clusters, methylation-based TF activity offers a more specific readout of TF-driven regulation of T-cell memory formation. In this process, FOXP1 and IRF4, known memory formation markers in T-cells, exhibited epigenetic concordance across modalities. In memory T-cells, IRF4 motifs displayed increased accessibility and decreased methylation compared to naive T-cells. In contrast, FOXP1 displayed the opposite pattern with limited changes, consistent with previous studies reporting accessibility changes in these TFs [33]. We found that AP-1 factors, such as BATF, exhibited starkly altered activity in memory formation, indicated by increased accessibility and decreased methylation. In contrast, NF-κB factors were predominantly influenced by methylation changes with comparatively minor accessibility shifts. This suggests that different TF families are governed by distinct epigenetic mechanisms that regulate T-cell memory. Further experimental and computational validation is needed to examine the interactions between these TFs, particularly how varying accessibility and methylation levels among different TF networks coordinate together to define memory cell identity. Our TF activity analysis also revealed that differences in accessibility between naive and CD8 memory T-cells are more pronounced than in CD4 memory T-cells (Figure 4D, Supplementary Figure 6A), consistent with the previous observations in the literature [40], indicating more substantial TF-associated chromatin remodeling. For example, EOMES exhibited increased accessibility with only a minimal decrease in DNA methylation in CD8 T cells, whereas CD4 memory T-cells showed only a limited change. However, since a portion of our data for matching multimodal single-cell epigenomes was obtained from our HIV exposure cohort, this may be the effect of HIV induced exhaustion of CD8 T memory cells and a general depletion of the CD4 population. Further experimentation is necessary to validate the TF binding dynamics of EOMES upon HIV exposure.

Consistent with our accessibility findings, the most extensive epigenetic remodeling in DNA methylation was observed in monocytes from severe COVID-19 cases compared to healthy controls. TF enrichment analysis in severe DMRs revealed a large overlap with the set of TF motifs enriched in accessibility analysis. This overlap suggests a coordinated remodeling of the TF-driven cytokine regulatory network involving both epigenetic modalities, rather than independent layers of regulation. Moreover, multimodal TF activity analysis confirmed that these TFs enriched in cCREs, such as AP-1 and NF-kB factors, were also differentially active, highlighting their role in severe COVID-19.

We further supported these findings through TF motif footprinting, which revealed that under severe COVID-19 conditions, monocyte regulatory programs undergo selective remodeling. Specifically, while generic AP-1 actors such as JUN displayed increased accessibility, specific inflammatory heterodimers like FOS::JUN exhibited hypermethylation in severe cases compared to controls. This implies that DNA methylation acts as a selective epigenetic constraint on inflammatory potential, even though the chromatin remains open. In contrast, NF-κB factors such as REL and RELA display a limited decrease in accessibility without accompanying methylation changes in TF footprints. The absence of accompanying DNA methylation changes in NF-κB sites suggests that they are likely not permanently silenced. This differs from lineage factors, which show a stable locked state due to hypermethylation. Consistent with the previous reports of Wimmers 2023 and Wilk 2021, our findings indicate stable activation of AP-1 factors in severe COVID-19 [5,26]. This implies that AP-1 remodeling may underlie dysregulated monocyte responses and sustained inflammation. Our results inferring TF activity of monocytes identified the pivotal AP-1 factors that are activated and NF-κB factors transiently modulated in severe COVID-19.

We acknowledge several limitations of our study. First, although our single-cell atlas profiled cohorts for influenza and organophosphate (OP) exposures, these conditions exhibited relatively limited differential chromatin accessibility at the cCRE level relative to variations observed across controls. We consequently prioritized analysis of the stronger epigenetic remodeling observed in COVID-19 and HIV exposures. However, even within this limited signal, we observed biological coherence. Our cross-exposure comparison analysis revealed that chromatin accessibility changes in Influenza exposures were weakly positively correlated with changes observed upon SARS-CoV-2 exposure, despite a lower magnitude, particularly in myeloid cells (Supplementary Figure 5). Secondly, although our motif-based inference of TF activity and footprinting infers TF binding dynamics from accessibility and methylation indirectly by aggregation across genome locations, it can prioritize TFs with potential driving roles of cell state regulation, a direct genome-wide quantification of TF binding could strengthen the results shown here.

In summary, our comprehensive multimodal atlas of human immune cell epigenomic remodeling across diverse environmental stimuli quantifies epigenetic signatures of T-cell exhaustion dynamics in chronic HIV infection, the regulatory TFs involved in T-cell memory formation, and epigenetic deregulation in monocytes in severe COVID-19 disease. By integrating chromatin accessibility with DNA methylation at single-cell resolution, this work establishes a framework for dissecting stimulus-specific transcriptional regulation, providing a valuable resource for future studies on immune response and dysfunctions associated with environmental exposures.

## Methods

### Sample acquisition and patient consent

Human peripheral blood mononuclear cells (PBMCs) were collected and isolated from different subjects in different cohorts exposed to various chemical or biological pathogens, including organophosphates, Influenza vaccines (followed by live Influenza virus challenge), HIV, and SARS-CoV-2 viruses. The subjects or surrogates have informed and approved all patient consents in different cores.

### snATAC-seq library generation, sequencing, and alignment

The snATAC-seq assay was performed by using the Chromium Next GEM Single Cell ATAC Reagent Kits version 1.1 (10x Genomics; PN-1000175) with the demonstrated protocol (nuclei isolation for scATAC-seq from PBMCs) provided by 10x Genomics. Briefly, cryopreserved PBMCs with around 100,000 cells were thawed and resuspended in RPMI medium, washed with PBS twice, and then lysed with 100 µl lysis buffer for 5 min. The lysed nuclei were centrifuged after washing with 1 ml washing buffer and then resuspended in a diluted nuclei buffer. The nuclei concentration was then determined with a TC20 Automated Cell Counter (Bio-Rad Laboratories; 1450102), and around 8,000-10,000 single nuclei were used for downstream steps: Tn5 transposition, single nuclei barcoding, and library preparation following the instructions in the kit. The final DNA libraries were sequenced with the NovaSeq 6000 system (Illumina), leading to around 200 million read pairs per sample. Data processing was performed with cellranger-atac software (10x Genomics; version 1.2.0). The demultiplexed sequencing reads were aligned to the GRCh38 (hg38) human reference genome, resulting in the related BAM files and fragment files for downstream analysis.

### snATAC-seq quality control, dimensionality reduction, clustering, and batch correction

Following preprocessing, we utilized the ArchR package (version 1.0.2) [40] to process snATAC-seq fragments. We added human reference genome (hg38) annotation with ArchR’s “addArchRGenome” function. Transcription start site (TSS) enrichment and the number of unique fragments were computed to assess the quality of the cells. Cells with TSS enrichment above 4 and log(nFragments) above 3 were retained. ArchR’s “addDoubletScores” was used to identify doublets, and “filterDoublets” with a ratio of 1.0 was used to remove doublets. 12,684 cells were identified and removed as doublets. 271,299 cells were retained after the initial filtering and quality control processes.

ArchR’s implementation of iterative LSI used dimensional reduction by “addIterativeLSI’’ with 25,000 variable features (tiling regions) and 30 dimensions. We identified clusters using “addClusters” with 0.8 resolution and obtained 22 clusters. For integration across exposure cohorts, Harmony batch correction was applied using “addHarmony” with “iterativeLSI” as reduced dimensions. Then, “addUMAP” was used to obtain the two-dimensional representation of the data with reducedDims = “Harmony”, nNeighbors = 40, minDist = 0.5, and metric = “cosine”. Batch-corrected data is only used for visualization purposes.

Gene activity score, weighted chromatin accessibility around each gene, of marker genes obtained using “getMarkerFeatures”. We then visualize these markers’ smoothened gene activity score on UMAP, where smoothening is applied using the MAGIC algorithm through ArchR’s “addImputeWeights**”**. Data smoothing was only used for visualization and not for downstream analyses. “addMotifAnnotations” adds the “JASPAR2020” motif set annotation. Deviation scores are computed using “addDeviationsMatrix”, and the deviation Z-scores of marker motifs were plotted on UMAP. Finally, track plots of marker genes are plotted using “plotBrowserTrack” from the ArchR package.

### Cell-type annotation, peak-calling, and generating pseudo bulk samples

Before peak calling, pseudo-bulk replicates are generated per cluster using “addGroupCoverages”. Then, we performed peak-calling with MACS2, using ArchR’s “addReproduciblePeakSet” with default settings. Canonical correlation analysis-based integration was performed to integrate the scRNA-seq ImmuneAtlas dataset [19] and scATAC-seq through ArchR’s “addGeneIntegrationMatrix” with default settings. We computed the Jaccard Index of scATAC cluster labels and their corresponding predicted scRNA labels. The final cell types were annotated to clusters using Jaccard similarity following integration (Supplementary Figure 1D). We assessed the significance of cell-type proportions by exposure condition as follows: The cell-type proportions per sample were obtained using the “prop.table” function. To compare disease versus control conditions, we generated boxplots with ggplot2 [42] and applied “stat_compare_means” from the ggpubr package [43] using the Wilcoxon rank-sum test and Bonferroni correction for multiple testing.

For pseudobulk analyses, we summarized fragment counts per cell type and sample as follows: Cell-type-specific fragments were extracted from Arrow files using “getFragmentsFromArrow”. Samples that contained fewer than 50 fragments per cell type were discarded from the analysis. In total, 688 pseudobulk samples were retained for analysis. Cell-type-specific fragments are then exported as bed files using “export.bed” from the rtracklayer R package [44].

TF motif cluster annotations [45] were added to the ArchR project using “addPeakAnnotations” from the ArchR package. Deviation scores for Viestra motifs were computed using “addDeviationsMatrix”. The added deviation Z-score matrix with annotated cell types was then extracted using the “getGroupSE” function with the following parameters: groupBy =” ClusterCellTypes”, divideN=TRUE.

### Cell-type-specific differential accessibility analysis using pseudo-bulk samples

Pseudo-bulk samples per cell type and sample were processed, and differential analysis was performed using the “run_chraccr” function from the ChrAccR package [46]. The configuration of vanilla ChrAccR analysis is as follows: differentialColumns =sample_exposure_group, filteringCovgCount=1L, filteringSexChroms=TRUE, filteringCovgReqSamples=0.005, differentialCutoffL2FC=0.5. By default, ChrAccR uses the DESeq2 package for differential analysis, which requires at least 3 samples per comparison group to perform pairwise differential analysis. For the COVID-19 cohort, the data processing date was used as the batch confounder by setting differentialAdjColumns = “processing_date” in the ChrAccR configuration. TF motif enrichment was done by LOLA implementation in ChrAccR by configuring the LOLA database to “lolaDbHg38”. Enrichment for TF motifs in differentially accessible regions for both hyper- and hypo-accessible regions was based on a Q-value threshold (q =< 0.05).

### Differential gene expression analysis of CD14+ Monocytes in COVID-19 using scRNA-seq and cross-modality comparisons

Samples from our COVID-19 cohort have paired publicly available single-cell transcriptomics data [3] for 34,548 cells. We subsetted CD14^+^ classical monocytes from the available Seurat object (see Supplementary Data) and then retained only protein-coding genes from the subsetted Seurat object. Next, we ran a pairwise differential analysis between the control and severe groups of COVID-19 with “DEseq2” using the “FindMarkers” function from Seurat [48]. Simultaneously, we subsetted CD14^+^ classical monocytes coming from the COVID-19 cohort with the same set of genes from the chromatin-derived gene-activity matrix in ArchR and ran the Wilcox test between the control and severe groups. We plotted a scatter plot of fold changes from the ArchR and Seurat analysis described above while highlighting the differential protein-coding genes (adj. p-value < 0.05).

### Processing and integrating snmC-seq with snATAC-seq

snmC-seq data for 38 samples and 22,488 cells corresponded to 110,376 cells from the scATAC-seq dataset. In total, 16,835 cell profiles were obtained for the COVID-19 cohort. A fluorescence-activated cell sorting (FACS) antibody cocktail was used to sort the immune cell types into 384-well plates based on their antibody labeling. Raw ALLC files from snmC-seq [16] were processed by summing CpG methylation, and total counts were over peak regions obtained from our scATAC dataset using ArchR’s “getPeakSet” function. Sex chromosomes and regions that weren’t covered by at least 20 percent of all cells were removed from the dataset. To obtain a feature matrix for integration with chromatin accessibility data, an inverted binarization procedure was applied to the retained DNA methylation dataset by setting entries in the cell-by-peak matrix with DNA methylation values below 0.4 to 1 and all other values to 0, for subsequent imputation. Missing values (NA) were preserved during this step and not coerced to 0, in order to allow for downstream imputation. Then, iterative LSI and clustering were applied to the binarized dataset with the following parameters: 50 dimensions, imputeMode=”pcimpute”, and 20,000 features. The details of the imputation process are described below. Reduced dimensions were then visualized on UMAP using the uwot package [47]. Imputed data is only used for iterative LSI and clustering, not for downstream processes.

We imputed missing values in the sparse DNA methylation matrix as follows: in the first iteration, the input methylation matrix is centered (but not scaled), and initially, each missing value in the input matrix is replaced with zero. Then PCA is performed using the irlba package [49]. The matrix is then reconstructed from the principal components by multiplying the scores (PCA-transformed values) with the transposed loadings (eigenvectors). This reconstructed matrix provides updated estimates for the missing values, which are then used to replace the initially imputed zeros. This process is repeated iteratively, updating the missing values with the estimated ones in each iteration until convergence. The convergence criterion is based on the mean squared difference between consecutive imputations and stops when the estimated imputation values change insignificantly. This procedure is iteratively performed until convergence. The resulting imputed data is only used for visualization purposes. To integrate chromatin accessibility data with the DNA methylation dataset from matched samples, the chromatin accessibility data were binarized by setting the regions with at least one Tn5 insertion and the rest to 0. Term frequency-inverse document frequency (TF-IDF) transformation was applied to binarized chromatin accessibility and inverted-binarized DNA methylation data of the shared samples. The resulting matrices were then used to create Seurat objects [48]. Canonical correlation analysis (CCA)-based integration is performed via “FindTransferAnchors” from the Seurat package with default parameters. We then computed nearest neighbors in CCA embeddings for both datasets and visualized the mapped cell labels on UMAP for each dataset.

### Cell-type-specific differential methylation analysis using pseudo-bulk samples

The custom function “createPseudoBulks’’ (see Supplementary Code) generates pseudo bulk samples by summing the methylated and unmethylated counts per site across the donors, cell type, and exposure condition. For each site, the methylation level is subsequently computed as the ratio of methylated counts to the total number of reads (the sum of methylated and unmethylated counts). It takes the maximum single-base coverage (default: 99999), sample annotation, grouping variables for summarizing methylated sites, excluded chromosomes (default: sex chromosomes), and strand specificity (default: FALSE) as parameters. The resulting pseudo-bulks were saved as bed files. In total, 1015 pseudobulk samples were generated.

For the analysis of methylation pseudobulk samples, the RnBeads package version 2.12.2 is used with the following settings: region.aggregation = “sum”, filtering.sex.chromosomes.removal = TRUE, and strand.specific = FALSE [50]. scATAC-seq peak regions were used as the reference regions for aggregating methylation counts. Differential analysis was conducted with RnBeads’ “rnb.run.differential” wrapper function for limma-based differential analysis [51], where each exposure group was compared with the control samples per cell type. LOLA enrichment analysis was performed using the same settings as described in the ATAC-seq analysis.

### Defining TF regulator activity from DNA methylation and cross-modality TF activity analysis

We computed a TF activity score based on how the DNA methylation signal deviates at the motif centers from the genomic background computed at +/-250bp distance away from the motif centers (Supplementary Figure 6A). To account for potential bias incurred by DNA sequence composition, we calculated both an observed and an expected deviation score and subtracted the expected deviation score from the observed one. The observed deviation score is computed for each sample by aggregating the methylated and unmethylated counts across putative binding sites of TFs (TFBS) and their flanking regions to determine the methylation percentage at each site. The methylation percentage is calculated as the ratio of methylated counts to the total read depth (the sum of methylated and unmethylated counts) for each sample. The final deviation score is then derived as Deviation score = (Average methylation in footprint base) / (Average methylation in the background).

To compute the expected methylation levels, we first divided the whole human genome into 30 base-paired bins and subsetted the distal enhancer regions [51]. We then calculated the GC frequency for each bin and discretized these into five frequency levels. For each frequency level, we computed the average methylation level across corresponding bins, which we refer to as the GC frequency table. These average values were then used to compute the methylation levels of the corresponding bins around TFBS in each sample and for each TF, which are referred to as the expected average methylation levels. We computed the expected deviation score using these expected methylation levels by dividing the expected methylation level at the motif centers by the average of the expected methylation levels at the motif flanking regions. We subtracted the expected deviation score from the observed deviation score to obtain the bias-corrected deviation score.

For heatmap visualizations, we applied row-wise Z-score transformation to normalize the bias-corrected TF deviation scores per TF motif. We developed an R package, methylTFR, and methylTFRAnnotationCreator to derive transcription factor (TF) activity from DNA methylation and generate the annotations described above (See Supplementary Code). We presented pre-computed annotations for JASPAR2020 and Altius TF motif sets [31,45] for Human Genome assembly version 38 as an R package called methylTFRAnnotationHg38 (see Supplementary Code).

We used the “run_methyltfr” function from the methylTFR package to compute bias-corrected deviation scores from the DNA methylation dataset for the JASPAR2020 motif sets. This function takes a sample annotation, a genome-wide GC distribution object, transcription factor binding sites, an enhancer region, a minimum coverage threshold per site (default=5), and a precomputed GC frequency table per annotation as input parameters. To obtain bias-corrected TF deviation scores from the chromatin accessibility dataset for JASPAR2020 motifs, we used the chromVAR package [21].

We used chromVAR’s “differentialDeviations” function with the Wilcox test to identify differentially active TF motifs based on chromatin accessibility-derived TF deviation scores, using the parameters parametric=TRUE and alternative=”two.sided”. Additionally, we adapted this function to analyze DNA methylation-derived TF deviation scores. Motifs with adjusted p-values lower than 0.05 were saved as tsv files. The correlation between the chromVAR and methylTFR deviation scores among the shared samples was assessed using a Pearson correlation test implemented through the ‘cor’ function from R. We obtained the motif footprints for accessibility data using the “getFootprints” function from the ArchR package using the default parameters. To plot accessibility footprints, we used “plotFootprints” from ArchR with normMethod =”divide” and used the default parameters otherwise. For DNA methylation footprints, we used the ggplot2 package to plot motif footprints we implemented in the methylTFR package (see Supplementary Code).

## Resource Availability

Processed data is available on the Gene Expression Omnibus (GEO) under accession number GSE306525. All code to analyze the datasets in this manuscript can be found on GitHub (https://github.com/EpigenomeInformatics/exposure_atlas_manuscript). The R package to compute TF activity from DNA methylation can be found on GitHub (https://github.com/EpigenomeInformatics/methylTFR). Fully processed scRNA data from the COVID-19 cohort were downloaded from https://www.covid19cellatlas.org/index.patient.html.

## Author contributions

W.J.G. and F.M. designed the study. W.J.G., J.R.E., and F.M. provided supervision. B.W. and D.C.C. conducted the experiments and generated the scATAC-seq dataset. A.B., C.B., R.C., and J.N. performed snmC-seq2, with A.B., C.B., and R.C. contributing to the development of the PBMC gating strategy and FACS. I.B.G. and B.W. carried out the data analysis and generated the figures. M.H. and W.W. were involved in methylation data analysis. I.B.G., B.W., F.M., and W.J.G. wrote the manuscript. All authors read the manuscript and provided comments.

## Acknowledgments

We thank all members of the Ecker, Müller, and Greenleaf labs for helpful discussions and advice. F.M. was supported by a NanoBioMed Young Investigator Grant at Saarland University. Figure 1A was created using BioRender. This work was supported by the Defense Advanced Research Projects Agency (DARPA) through the DARPA Epigenetic Characterization and Observation (ECHO) program for the project Single-cell Analysis for Forensic Epigenetics (SAFE), issued by the US Army Research Office under cooperative agreement W911NF-19-2-0185. J.R.E. is an Investigator of the Howard Hughes Medical Institute. We would like to thank former DARPA Biological Technologies Office Program Manager Eric Van Gieson, as well as the current leadership team, including Jean-Paul Chretien and Thomas Thomou, for their guidance and insightful comments. W.J.G. acknowledges funding support from National Institutes of Health (NIH) grants P50-HG007735 and UM1-HG009442, UM1-HG009436. V.G.F acknowledges funding from NIH grant 1R01AI165671. T.E. acknowledges Armen Donabedian. We would like to thank all the unidentified donors who contributed biological samples for this project through our collaborators.

---

**Supplementary Figure 1:**
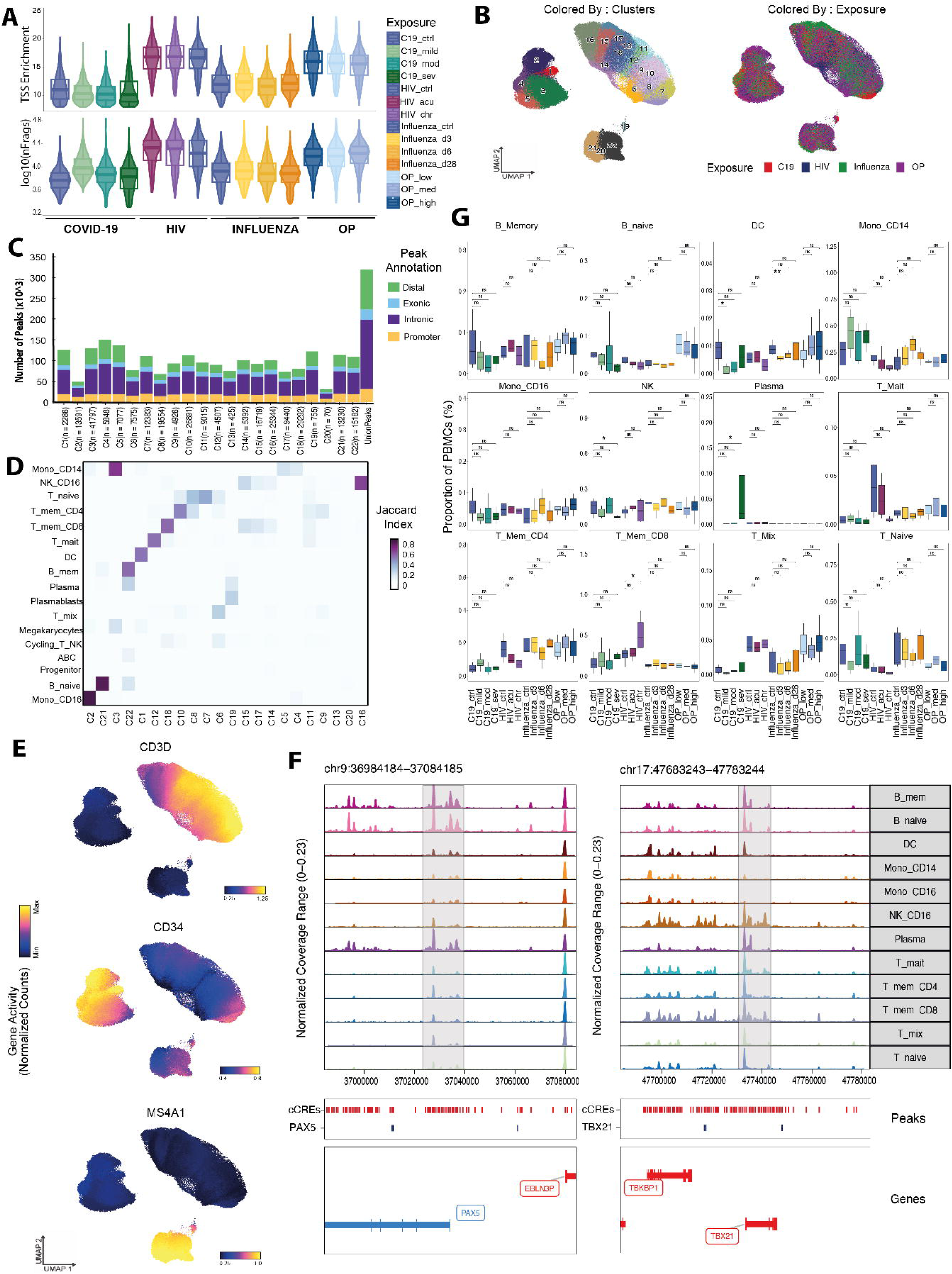
Quality control, cell-type annotation, and TF motif variability in the single-cell ATAC dataset. A) Violin plot of transcription start site (TSS) enrichment (top) and the number of unique fragments (bottom) across the exposure conditions B) UMAP projections of full scATAC dataset, colored by Louvain clusters (left) and colored by exposure cohort (right) C) Bar plot of the number of peaks by the number of unique fragments of each cluster, colored by peak annotation D) Correspondence between scATAC clusters and scRNA labels colored by Jaccard Index E) UMAP projections of the full scATAC dataset colored by relative gene activity scores of canonical markers (CD3D, CD34, and MS4A1) F) Genome accessibility tracks by cell type with a focus on genes (PAX1 and TBX21) and, their respective motif occurrences

**Supplementary Figure 2:**
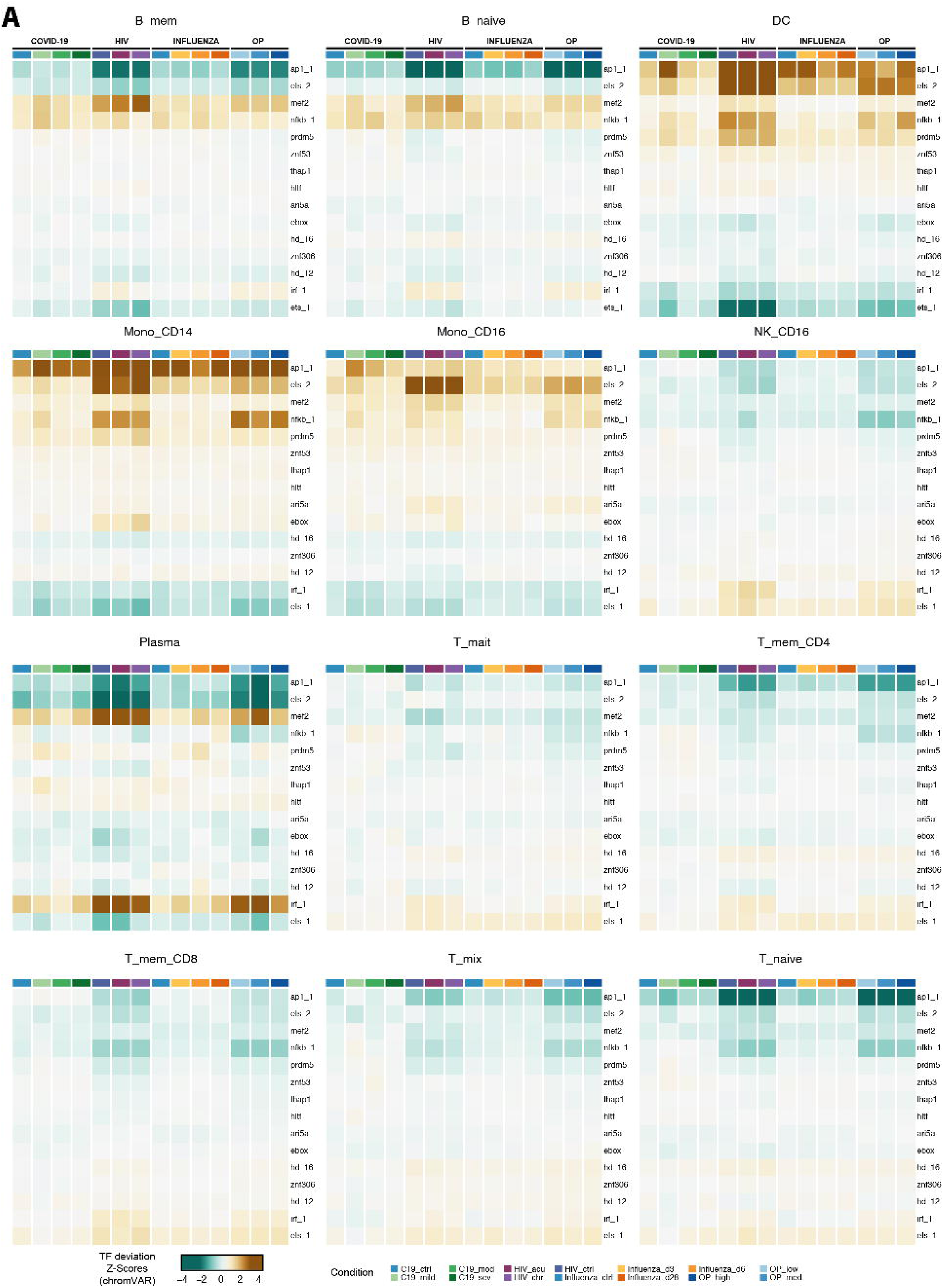
Transcription factor motif accessibility profiles across immune cell subsets and exposure conditions. A) TF motif accessibility by cohort and cell type. Heatmaps display chromVAR deviation Z-scores representing TF clusters [45]. Rows denote TF clusters with representative factors: ap_1 (FOS/JUN), nfkb_1 (REL/NFKB1), irf_1 (IRF1/7/8), and ets_2 (ETS2/FLI).

**Supplementary Figure 3:**
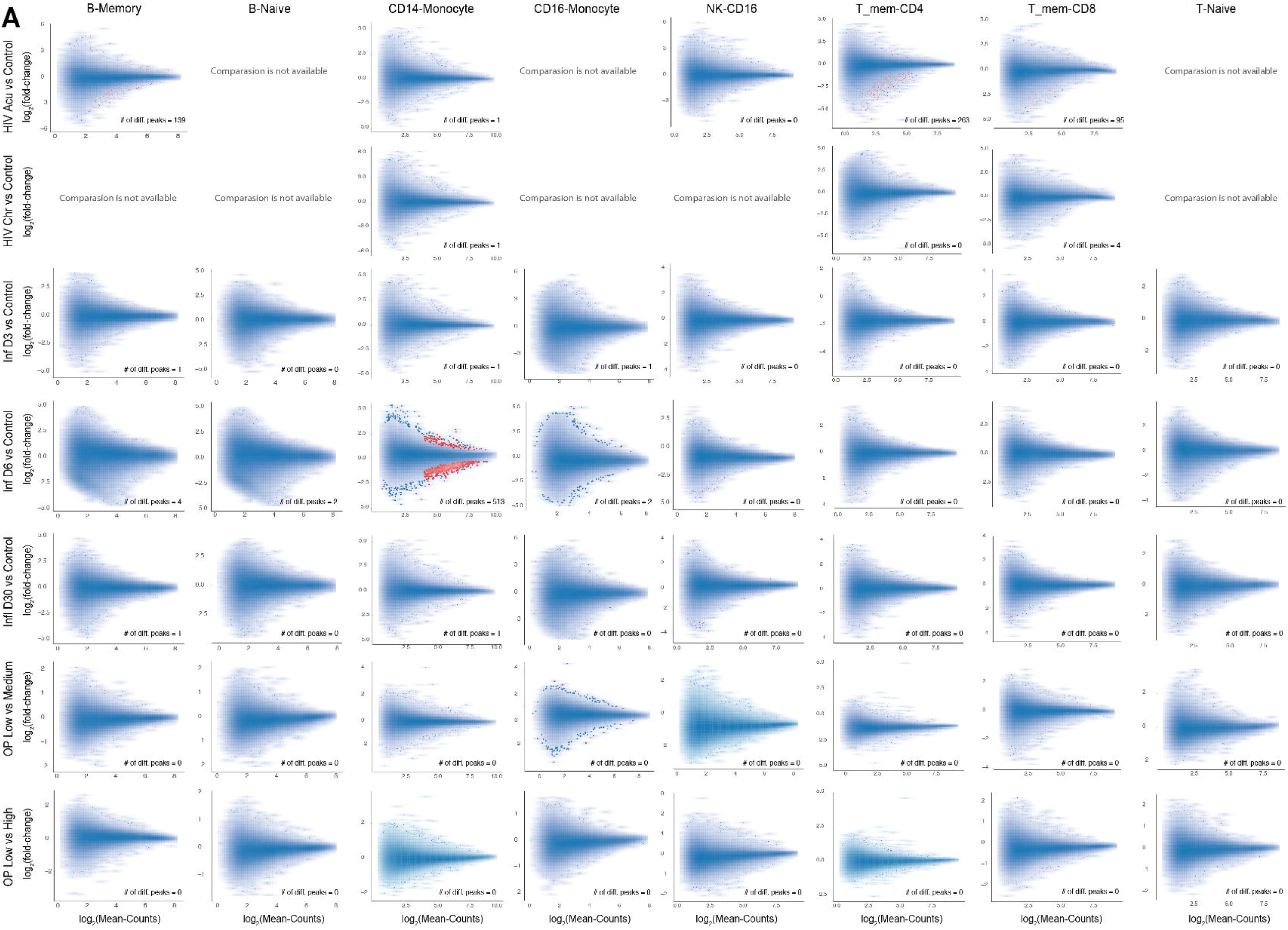
Differential chromatin accessibility across exposure conditions. A) MA plots of cCREs per cell type and exposure condition comparisons; Comparisons with insufficient sample size for differential analysis are marked “Comparison is not available.” Significant DARs are highlighted with red color (p.adj < 0.05 and |log2(fold-change) >= 0.5) |. Point density is visualized using 2D kernel density estimation; outlier points from low-density regions (<0.1% of all points) are plotted individually.

**Supplementary Figure 4:**
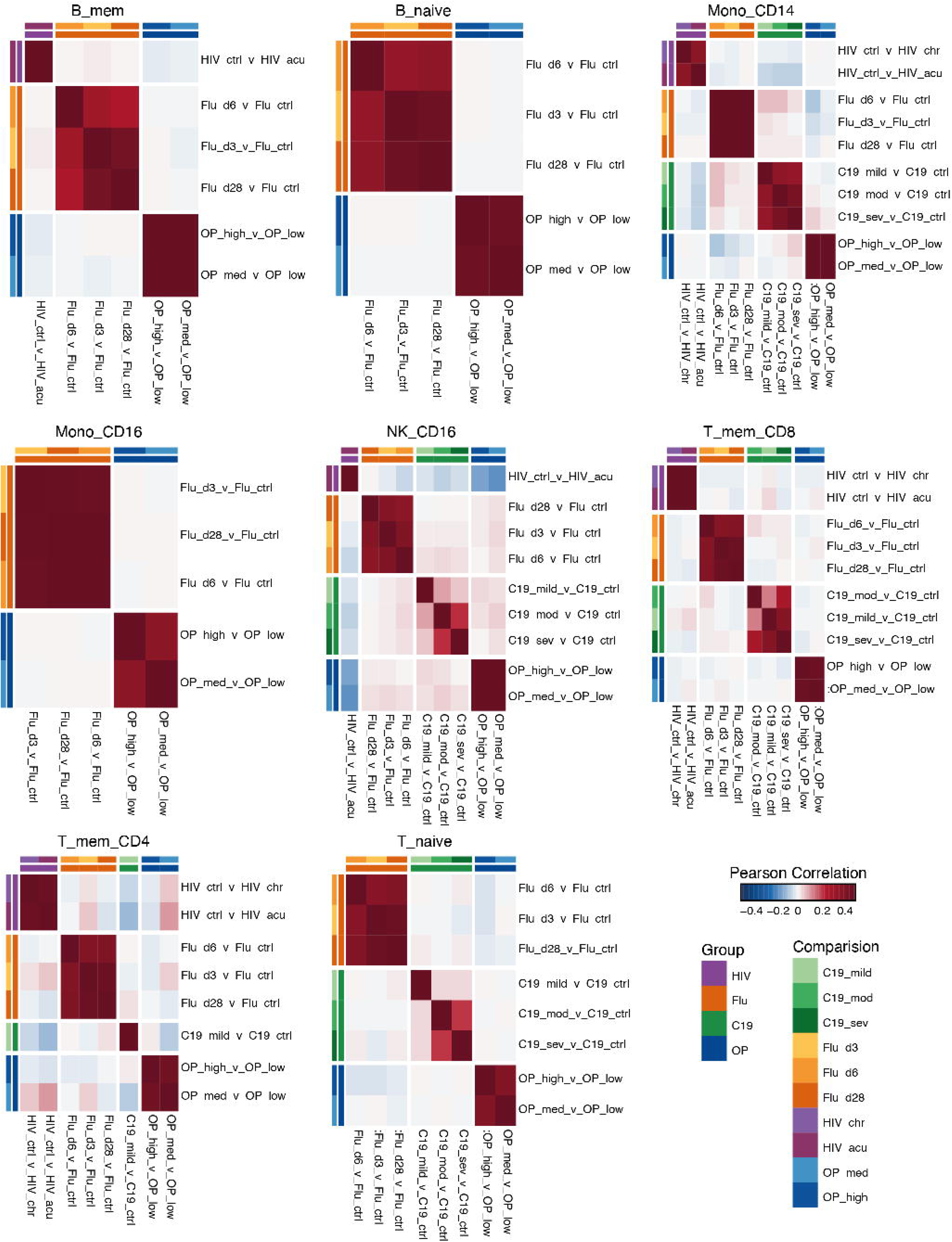
Pairwise correlation analysis of chromatin accessibility changes across viral and chemical exposures. A) Pairwise Pearson correlation of fold-change profiles across pairwise comparisons of all exposure conditions to respective control groups. Rows and columns are annotated by cell type, comparison group, and exposure. Heatmap colors indicate correlation strength, where blue is negative correlation, white is near zero, and red is positive. Comparisons in cell types with insufficient cell numbers for a differential accessibility analysis have been omitted.

**Supplementary Figure 5:**
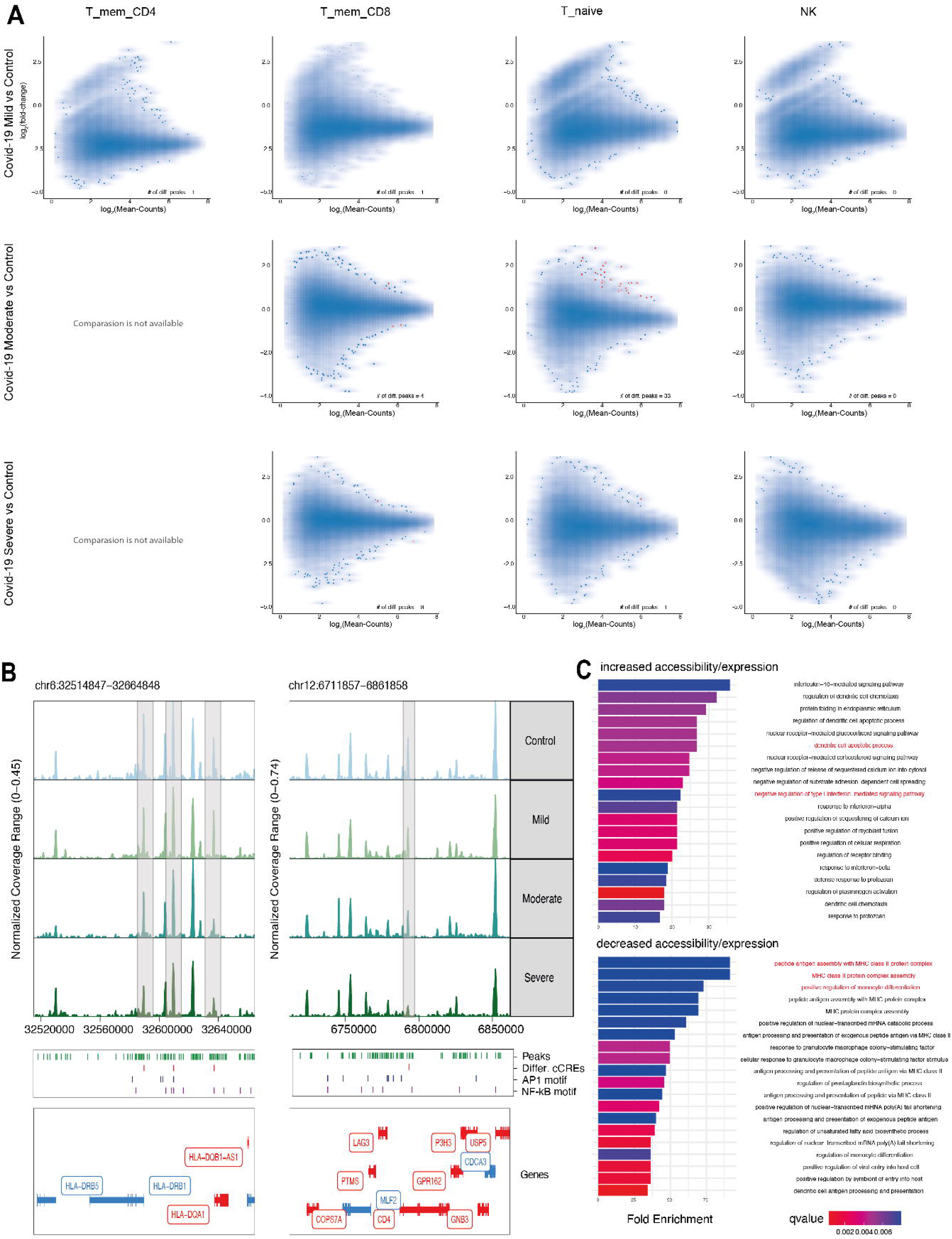
Differential accessibility analysis in the COVID-19 cohort. A) MA plots of cCREs per cell type and COVID-19 condition comparisons; Comparisons with insufficient sample size for differential analysis are marked “Comparison is not available”. Significant DARs (adj.p < 0.05 and |log_₂_FC| ≥ 0.5) are shown in red. Point density is visualized using 2D kernel density estimation; outlier points from low-density regions (<0.1% of of all points) are plotted individually. B) Genome accessibility track visualization of selected DARs upon SARS-CoV-2 exposure with highlighted genes, including HLA Class II families, and CD4, with AP-1 and NF-kB motif loci highlighted. C) Fold enrichment analysis (fold enrichment, colored by qValue) results for genes with increased expression/accessibility (top) and genes with decreased expression/accessibility (bottom), with previously reported genes highlighted in red [53]

**Supplementary Figure 6:**
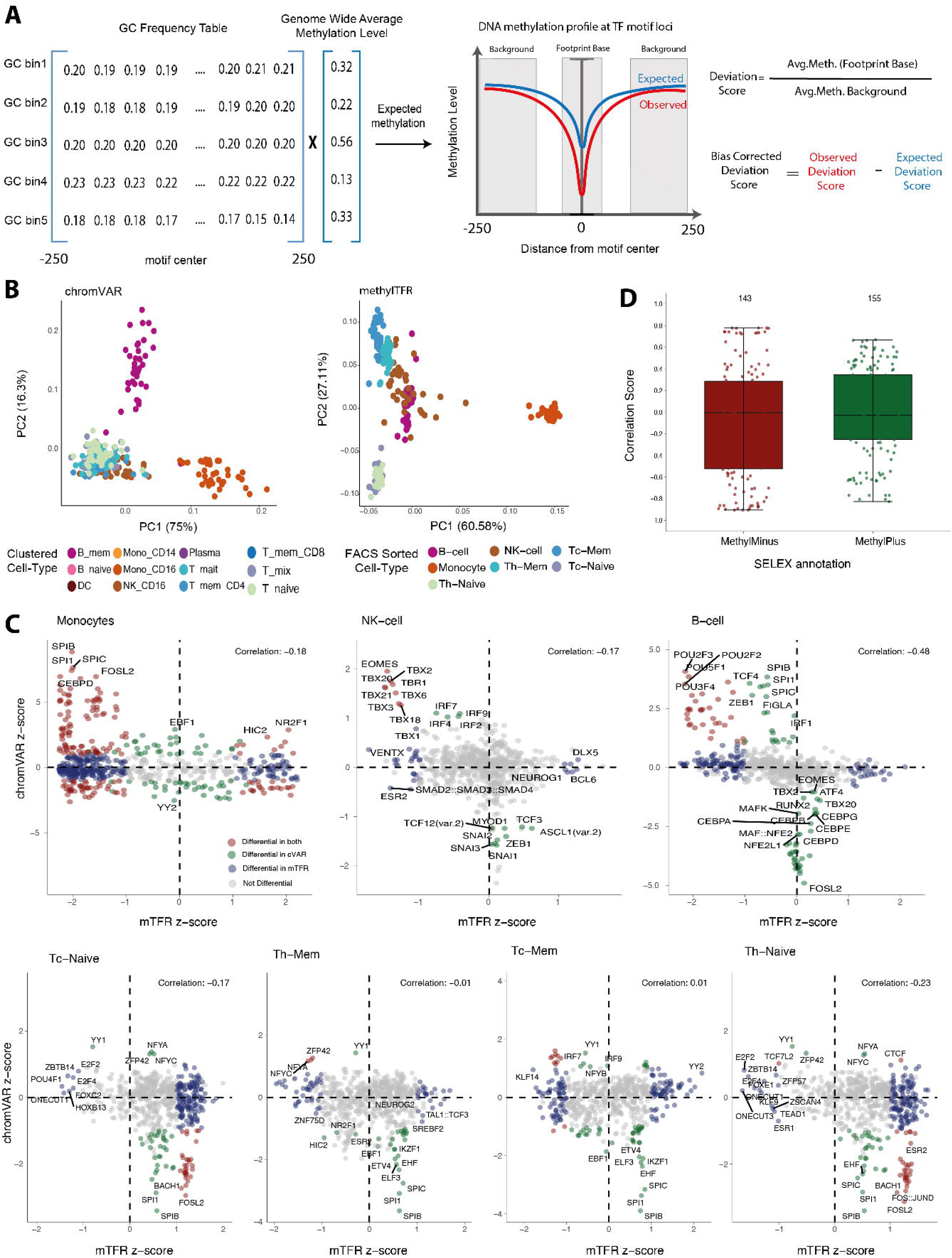
Multimodal TF activity analysis across cell types. A) Schematic of motif footprint-based TF deviation score computation with methylTFR, where expected methylation levels are represented as blue, while the observed levels are represented as red. B) PCA plots of chromVAR deviations (left) with snmC-mapped cell types and methylTFR deviations (right) with sorted cell types. C) Boxplots of Spearman correlation coefficients between chromVAR and methylTFR TF activities across methyl-SELEX labels, with the number of motifs per category annotated on top. D) Scatterplots comparing motif activity per cell type. Motif activity was defined as the z-score of methylTFR vs chromVAR on pseudobulk samples. Differentially active motifs in the two modalities (One vs other cell types, padj < 0.05) are highlighted as: red differential in both, green differential only in chromVAR, blue: only differential in methylTFR, gray: not-differential.

**Supplementary Figure 7:**
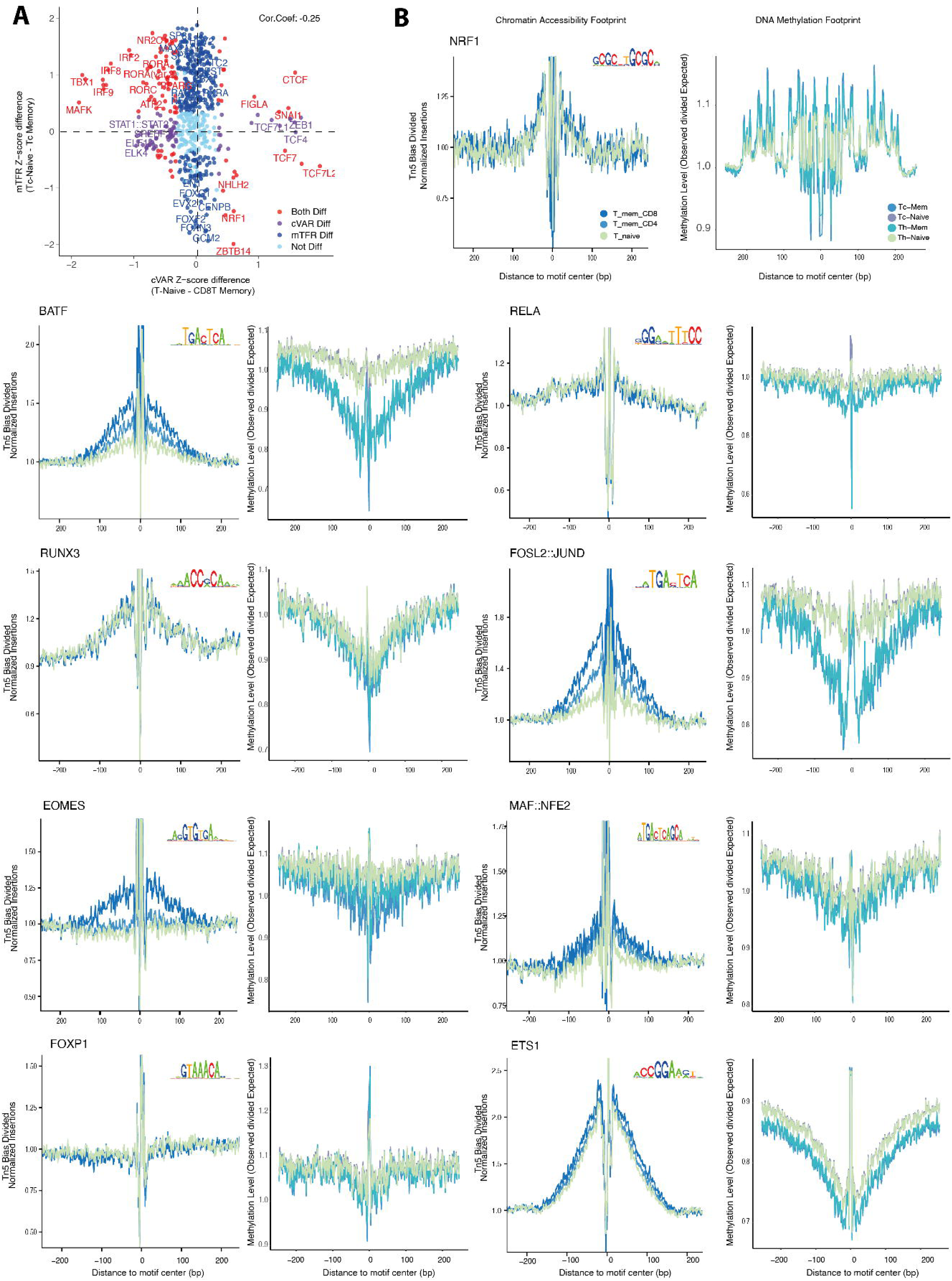
TF signatures of memory formation in T-cells. A) Z-score difference (CD8T-Memory Z-score subtracted from T-naive Z-score) between DNA methylation-based TF activity and chromatin accessibility-based TF activity, with highlighted differential motifs (Wilcoxon test, padj < 0.05). B) Side-by-side chromatin accessibility and methylation footprint plots of TF motifs with variable TF activity (NRF1, BATF, RELA, RUNX3, FOSL2::JUND, EOMES, MAF::NFE2, FOXP1, and ETS1, respectively) in naïve and memory T cell populations.

**Supplementary Figure 8:**
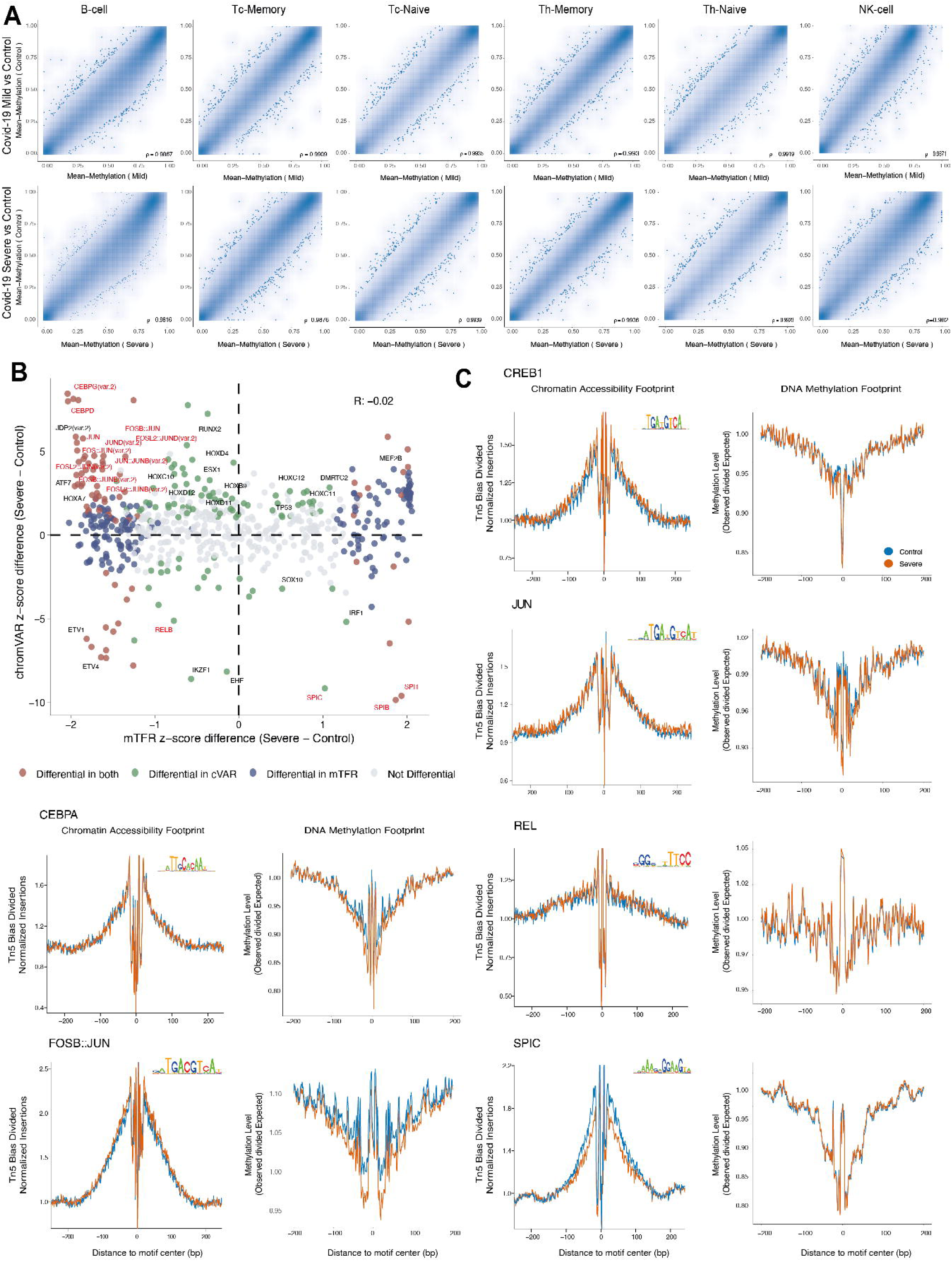
Differential DNA methylation analysis in the COVID-19 cohort. B) Density scatter plots of the DNA methylation levels in cCREs in the COVID-19 condition compared with controls per cell type. Point density is visualized using 2D kernel density estimation; outlier points from low-density regions (<0.1% of all points) are plotted individually. Scatterplot comparing motif activity in monocytes (severe mean Z-score - control mean Z-score). Motif activity was defined as the z-score of methylTFR vs chromVAR on pseudobulk samples. Differentially active motifs in the two modalities (severe vs control, padj < 0.05) are highlighted as: red differential in both, green differential only in chromVAR, blue: only differential in methylTFR, gray: not-differential. C) Side-by-side accessibility and methylation footprint plots of differentially active TF motifs (CREB1, CEBPA, JUN, REL, SPIC, and FOSB::JUN, respectively)

